# Stable *C. elegans* chromatin domains separate broadly expressed and developmentally regulated genes

**DOI:** 10.1101/063990

**Authors:** Kenneth J. Evans, Ni Huang, Przemyslaw Stempor, Michael A. Chesney, Thomas A. Down, Julie Ahringer

**Affiliations:** The Gurdon Institute and Department of Genetics, University of Cambridge, UK

## Abstract

Eukaryotic genomes are organized into domains of differing structure and activity. There is evidence that the domain organization of the genome regulates its activity, yet our understanding of domain properties and the factors that influence their formation is poor. Here we use chromatin state analyses in early embryos and L3 larvae to investigate genome domain organization and its regulation in *C. elegans*. At both stages we find that the genome is organized into extended chromatin domains of high or low gene activity defined by different subsets of states, and enriched for H3K36me3 or H3K27me3 respectively. The border regions between domains contain large intergenic regions and a high density of transcription factor binding, suggesting a role for transcription regulation in separating chromatin domains. Despite the differences in cell types, overall domain organization is remarkably similar in early embryos and L3 larvae, with conservation of 85% of domain border positions. Most genes in high activity domains are expressed in the germ line and broadly across cell types, whereas low activity domains are enriched for genes that are developmentally regulated. We find that domains are regulated by the germ line H3K36 methyltransferase MES-4 and that border regions show striking remodeling of H3K27me1, supporting roles for H3K36 and H3K27 methylation in regulating domain structure. Our analyses of *C. elegans* chromatin domain structure show that genes are organized by type into domains that have differing modes of regulation.

**Significance statement:** Genomes are organized into domains of different structure and activity, yet our understanding of their formation and regulation is poor. We show that *C. elegans* chromatin domain organization in early embryos and L3 larvae is remarkably similar despite the two developmental stages containing very different cell types. Chromatin domains separate genes into those with stable versus developmentally regulated expression. Analyses of chromatin domain structure suggest that transcription regulation and germ line chromatin regulation play roles in separating chromatin domains. Our results further our understanding of genome domain organization.

## INTRODUCTION

The complete genome sequence, which provides the information necessary for constructing an organism, is interpreted in the context of chromatin. Covalent modifications of histone tails and histone variants can regulate and/or reflect genome function, and so are markers of chromatin state and genomic activity (1). For example H3K4me3 often marks active promoters, H3K36me3 transcription elongation, and H3K27me3 Polycomb silenced regions. Previous studies showed that genomic regions of similar activity harbor shared combinations of modifications, termed chromatin states, and that subdividing the genome according to these combinations is a powerful method for annotation and uncovering novel functional regions (2–5). Here we apply chromatin state mapping to two developmental stages of the model organism *C. elegans* and use the resulting maps to investigate genome domain organization and its regulation.

*C. elegans* is highly amenable for global studies of chromatin structure and function because it has a small, well-annotated genome (30X smaller than human), and work of the modENCODE consortium has provided a large number of datasets mapping the locations of chromatin associated factors such as histone modifications and transcription factors (6–10). *C. elegans* chromatin shows features in common with those of other organisms, such as the type of marking at regulatory regions and at active or inactive genes (7, 10–12). Additionally, the derivation of a single set of chromatin states for *C. elegans*, Drosophila, and human using a single joint genome analysis and data from eight histone modifications highlighted the common properties of chromatin in the three organisms (10). However, because chromatin differences also exist, the jointly derived chromatin states are not suitable for *C. elegans* specific analyses, and no other *C. elegans* chromatin state maps have been published.

Previous studies have described broad properties of *C. elegans* genome organization. The distal “arms” and central regions of the autosomal chromosomes show differences in transcription activity, chromatin composition, and recombination rate (6, 7, 11, 13, 14). Central regions have higher average gene expression, moderate enrichment of histone modifications associated with active transcription, and lower meiotic recombination than distal arm regions. In contrast, most features associated with heterochromatin, such as H3K9 methylation and nuclear envelope association, are found on the chromosome arms (7, 15). However, the chromosome arms are not purely heterochromatic. Actively transcribed genes reside on the chromosome arms and these genes are marked by histone modifications associated with gene activity, as in the central regions (7). In addition, the X chromosome shows extensive chromatin differences compared to autosomes due to dosage compensation (16). These previous studies have provided a large-scale picture of *C. elegans* chromosome organization.

Here we investigate *C. elegans* chromatin and genome organization and its regulation through the generation and analyses of *C. elegans* specific chromatin state maps for early embryos and third larval stages. As in other organisms, chromatin states correlate with many biological features including enhancers, promoters, transcription elongation, gene ends, repeat regions, and inactive genes. Analyzing patterns of states revealed that chromatin domains of differing activity separate germ line and broadly expressed genes from developmentally regulated genes. The properties of domains and the border regions between them suggest that transcription regulatory regions and germ line chromatin marking play roles in domain separation. Our results provide a framework for future studies of chromatin structure and function in *C. elegans*.

## Results

### 20 state models of *C. elegans* chromatin

To investigate features and domain organization of *C. elegans* chromatin, we derived 20 state early embryo (EE) and third larval stage (L3) chromatin state maps, using Hidden Markov Models and chromatin immunoprecipitation data for 17 histones or modifications (see Methods). Patterns and levels of enrichment of many histone modifications differ on *C. elegans* autosomes compared to the X chromosome, reflecting dosage compensation (7, 16–18) which caused whole genome chromatin state maps to subdivide into separate autosomal and X-chromosome-specific states. Therefore, for each stage we generated a separate map for autosomes and the X chromosome (Fig. 1A, Fig. S1, Dataset S1). The EE and L3 autosomal chromatin states show much greater similarity to each other than do the EE and L3 chromosome X states (Fig. S1, S2), consistent with alterations in chromatin structure and marking induced by dosage compensation after the EE stage (16). The chromatin states were annotated by analysing the associations of states with a range of different genomic features (Fig. 1, S1, S3-S7). As well as differences in enrichment levels for histone modifications, the states show differences in median length (250-1250bp), genomic coverage (2.2-9%), and GC content (25-44%) (Fig S1). Below, we briefly describe chromatin state annotation and properties using L3 autosomes as an example. We then focus use the states to investigate autosomal chromatin domain organization and its regulation.

**Figure 1.**
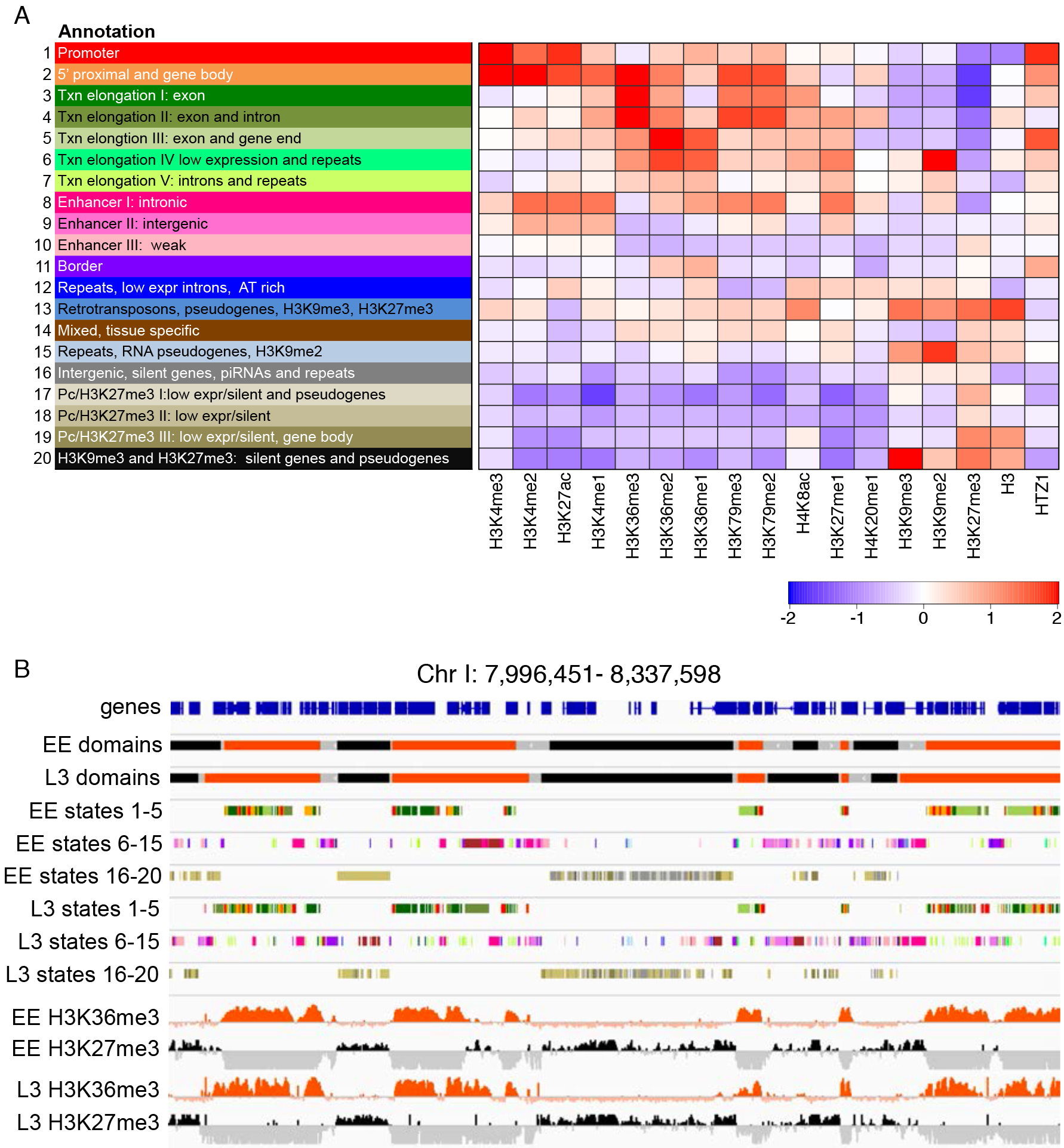
Chromatin states and domains. (A) L3 state key and annotation. Left panel gives state numbers and annotations; right panel shows relative enrichment or underenrichment of the indicated histones or histone modifications in each state. The scale bar shows the average z-score of the mark. (B) IGV screenshot of 340kb region on chromosome I (I:7,996,451-8,337,598) showing genes, domains, chromatin states, H3K36me3, and H3K27me3 in EE and L3.

### States associated with active genes and enhancers

We found that states 1-8 predominantly mark high and moderately expressed genes, with states 1-5 most associated with genes in the highest quintile of expression (Figs. S3, S5A). These chromatin states mark different types of genic regions: promoter (state 1), 5’ proximal (state 2), transcription elongation (states 3-7), and enhancer (state 8) (Figs. S1, S5A,B). The different transcription elongation states are associated with different expression levels and genic regions. For example, state 3 marks highly expressed exons whereas state 6 marks transcribed regions of lower expression, and state 5 typically marks gene ends (Fig. S3, S5A).

We define states 8-10 as likely enhancer regions based on their enrichment for chromatin modifications typical of enhancers (high H3K4me1, high H3K27Ac, low H3K4me3) and their association with annotated enhancers (Figs. S1, S5B). Annotated non-coding RNA genes that are unclassified in Wormbase are also frequently associated with states 8-10 suggesting that some of these may be enhancer transcripts (Fig. S5C). Other non-coding RNA genes (e.g, miRNAs, tRNAs, piRNAs) show different state enrichments (Fig. S5C).

### States associated with inactive genes

We found that inactive and lowly expressed genes are associated with states 16-20 (Figs.S3, S5A). In contrast to active states, inactive states usually do not mark particular gene regions but instead are more uniformly distributed across genes (Figs. S3, S5A). However, inactive states do show differential enrichment in genic versus intergenic regions (Fig. S3). Consistent with known associations of histone modifications with silenced genes (61), inactive states are enriched for H3K27me3 (a mark of Polycomb mediated silencing; states 17-19) or both H3K27me3 and H3K9me3 (a mark of heterochromatin; states 16 and 20) (Fig. S1). Co-occurrence of H3K27me3 and H3K9me3 in *C. elegans* has been noted previously (10).

Inactive states 16-20 show different chromosomal distribution patterns and genic associations (Fig. S5F). For example, inactive state 17, which is enriched for H3K27me3, is highly prevalent on chromosome V (Fig. S5F). Chr V is unique in harboring a large fraction (68%) of the 1383 odorant receptors annotated in *C. elegans* (50). These receptors are transcriptionally inactive in most cells, usually being expressed in only one or a few neurons (62). We found that state 17 is highly associated with odorant receptor genes, marking 62% of them genome-wide. In addition, state 17 marks 57% of odorant receptor pseudogenes (out of 290) and 18% of pseudogenes of other classes (Fig. S5F). The finding of a chromatin state associated with both pseudogenes and a class of widely silenced genes (odorant receptors) suggests that that these loci may be repressed by a shared mechanism.

### Mixed states

Because the histone modification mapping was conducted in whole animals, it was expected that states associated with tissue specific gene expression might display enrichment for histone modifications of both active and inactive genes. Indeed, states 13 and 14 display H3K27me3 pattern, and genes marked by these states are enriched for having high gene expression combined with high H3K27me3 (Fig. S5E). States 13 and 14 are also enriched for tissue specific genes identified by gene expression profiling (Fig. S5E).

### Repeats

The *C. elegans* genome harbors about 100,000 annotated repeat elements, which fall into about 163 families (52). We found that six chromatin states are highly associated with repeats (states 6, 7, 12, 13, 15, and 16; Fig S5D). Different repeat classes and individual repeats are associated with different chromatin states (Fig. S5D). The variation of chromatin states on different repeat types may reflect differences in their regulation or function.

### Chromatin states on autosomes demarcate chromatin domains of different activities

We next used the chromatin states to investigate chromatin domain structure and its regulation. We focused on autosomes because the chromatin states are highly similar in the EE and L3 maps, facilitating comparative analyses. In browsing, we observed that states associated with the highest or lowest quintiles of gene expression (states 1-5 and states 16-20, respectively; Fig. S3) were located in extended genomic domains interspersed with the other 10 states (Fig. 1B). Based on these patterns, we defined highly active domains, lowly active domains, and border regions separating domains in EE and L3 autosomal chromatin (Fig. 1B, Dataset S2, see Methods). In brief, regions containing states 1-5 and the neutral states among them (states 6-15) without interruption by states 16-20 were defined as highly active domains, ending with an active state. Similarly, regions containing states 16-20 and the neutral states among them without interruption by states 1-5 were defined as lowly active domains, ending with an inactive state. The genomic regions between highly active and lowly active domains were defined as borders. Table S1 gives statistics on domain sizes and numbers of genes in domains. Domains are larger and contain more genes than expected by chance (Table S1; see Methods). For example, in EE, the median highly active domain is 13054 bp (4506 bp expected) and contains four genes (one expected) whereas the median lowly active domain is 23874 bp (13647 bp expected) and contains four genes (two expected). Figure S8 shows the distribution of the 20 chromatin states and Fig. S9 the distribution of histone modifications in the different regions; we note that chromatin state 11 is highly associated with domain borders in both EE and L3 domain maps.

Although the domains are defined based on the location of chromatin states associated with high and low gene expression, they are not uniform in activity. For example in L3 larvae, 20% of genes in the top quintile of expression lie in lowly active domains, and 11% of genes in the bottom 40% of expression lie in highly active domains. Based on the analyses below, we will refer to the highly active domains as “active,” the lowly active domains as “regulated,” and the regions in between as “border.” Across each autosome, the distribution of active and regulated domains is relatively uniform, although chromosomes vary in the relative proportions of active and regulated domains (Fig. S10).

### Chromatin domain structure of early embryos and L3 larvae is strikingly similar

To investigate the developmental regulation of chromatin domains, we compared the positions of domains and borders in EE and L3 larvae. These two samples represent very different populations of cells. The profiled early embryo samples contained undifferentiated cells undergoing cell division (1 - 300 cell stage embryos) whereas the L3 larval samples contained ~85% differentiated somatic cells and ~15% mitotic germ cells (7). Surprisingly, the autosomal chromatin domain structure of early embryos and L3 larvae is strikingly similar, with 85% of EE border regions overlapping an L3 border. Additionally, 91% of bases in active domains and 89% of bases in regulated domains are in common between EE and L3 stages. The overall consistency in domain structure between early embryos and L3 larvae suggests that mechanisms determining shared organization are largely independent of cell fate.

### Properties of domains and borders

We next investigated chromatin domain properties. As expected, RNA polymerase II levels sharply increase at the transitions from borders to active domains (Fig. 2). The transitions from regulated domains to border regions and from border regions to active domains have low levels of histone H3, indicative of nucleosome depletion, suggesting that these regions are more accessible than the neighboring chromatin (Fig. 2). Intriguingly, two families of repeat elements (CELE1 and CELE2) are particularly associated with borders (Fig. S11). Additionally, border regions are enriched for enhancer chromatin states and distal transcription factor binding sites, typical of enhancers, suggesting that these regions have transcription regulatory activity (Fig. 2).

**Figure 2.**
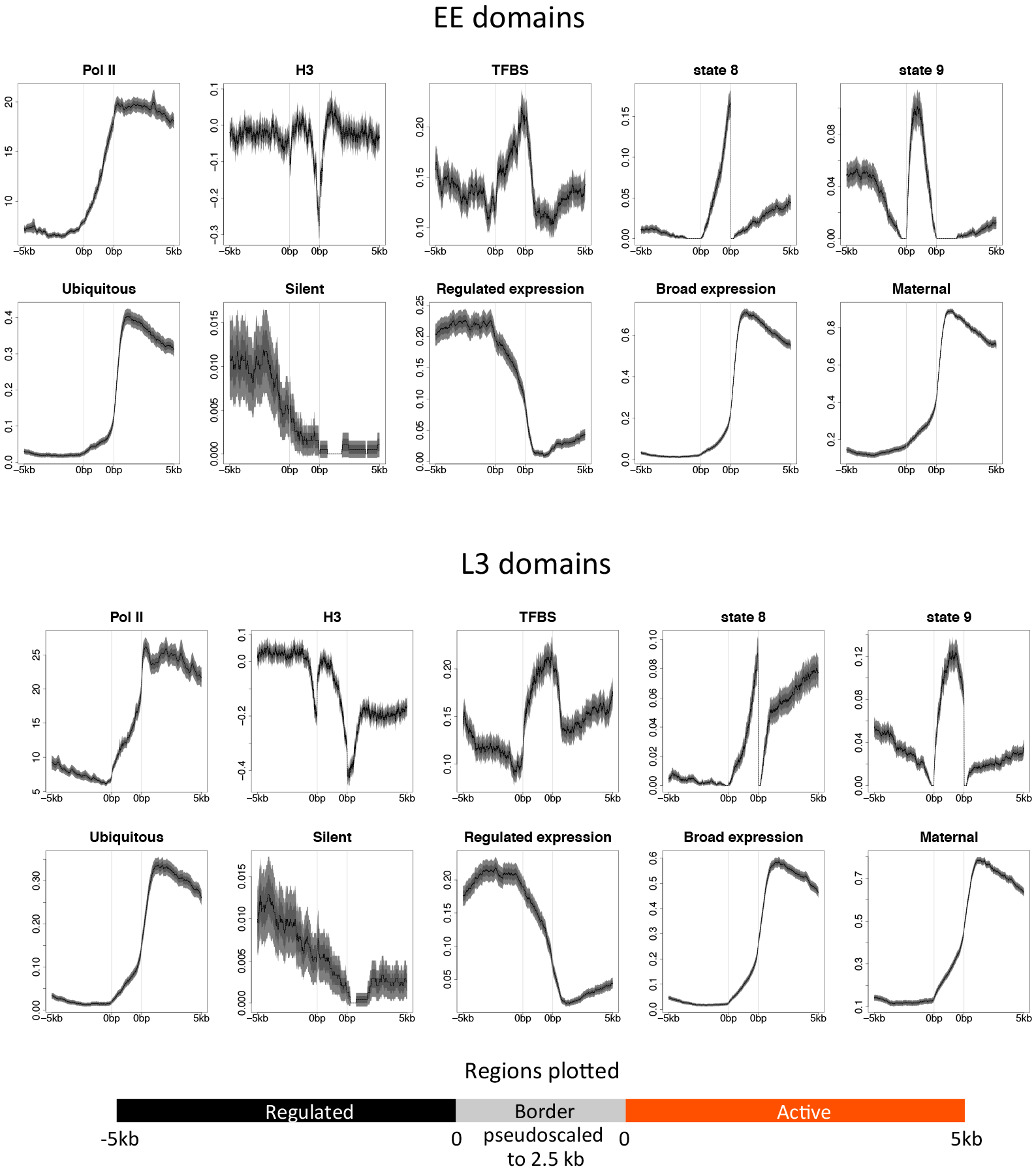
Properties of EE and L3 autosomal active, border, and regulated domains. Plots are centered and anchored at borders pseudoscaled at 2.5kb, and show 5 kb into regulated domains (left) and 5kb into active domains (right). Lines show mean signal, darker filled areas show standard error, and lighter filled areas are 95% confidence intervals. Grey vertical lines indicate edges of the border region.

The genes that reside in active and regulated domains have different properties. Not surprisingly, ubiquitously expressed genes lie predominantly in active domains, whereas silent genes (those with low or no detectable expression, dcpm <0.005 in all stages; n=764) are usually found in the lowly active regulated domains (Fig. 2). However, the majority of genes in regulated domains are detectably expressed at one or both stages (54% are in the top 60% of expression).

To further investigate the properties of genes in the different domains, we used the coefficient of variation (cv) of gene expression across 35 developmental stages and cell types (54). Genes expressed at a similar level across the 35 conditions (broad expression) have low cv values whereas genes differentially expressed across conditions (regulated genes) have high cv values. We considered genes in the bottom third of cv values as broadly expressed and genes in the top third as developmentally regulated. Gene expression variation (cv score) shows a remarkable association with domain type. Genes with broad expression across development and cell types (low cv) lie primarily in active domains whereas genes with developmentally regulated expression (high cv) are predominantly found in regulated domains (Fig. 2). Further, we found that most genes (86%) in active domains have maternally contributed mRNA indicating that they are expressed in the germ line (Fig. 2). We also observed that active domains contain high levels of H3K36me3 and that regulated domains contain high levels of H3K27me3 (Fig. 1B, 3A). Alternating blocks of H3K36me3 and H3K27me3 were previously noted in early embryo chromatin (19).

To summarize, the genome is organized into two types of domains: H3K36me3-rich “active” domains containing genes that are broadly and germ line expressed, and H3K27me3-rich “regulated” domains containing genes that have highly regulated or low expression. The correspondence between domain type and H3K36me3 or H3K27me3 levels suggests that these modifications may play roles in defining active and regulated domains.

### A role for MES-4 in domain definition

The association of active domains with maternal gene expression and high levels of H3K36me3 prompted us to investigate a possible relationship with MES-4, a germ line H3K36 histone methyltransferase. Two H3K36me3 methyltransferases have been studied in *C. elegans*. MET-1 encodes a Set2 family transcription coupled H3K36 methyltransferase active in most cells (20–22). MES-4 is a germline specific NSD family H3K36 histone methyltransferase with transcription independent activity (21–23). In the germ line, MES-4 marks expressed genes with H3K36me3, and this germ line marking is inherited and maintained in early embryos by maternally contributed MES-4 (21–23). Following knockdown of *mes-4*, early embryos show reduced H3K36me3 on germ line expressed genes, which is accompanied by increased H3K27me3 (19). As expected, MES-4 is enriched in active domains (Fig. 3A)

**Figure 3.**
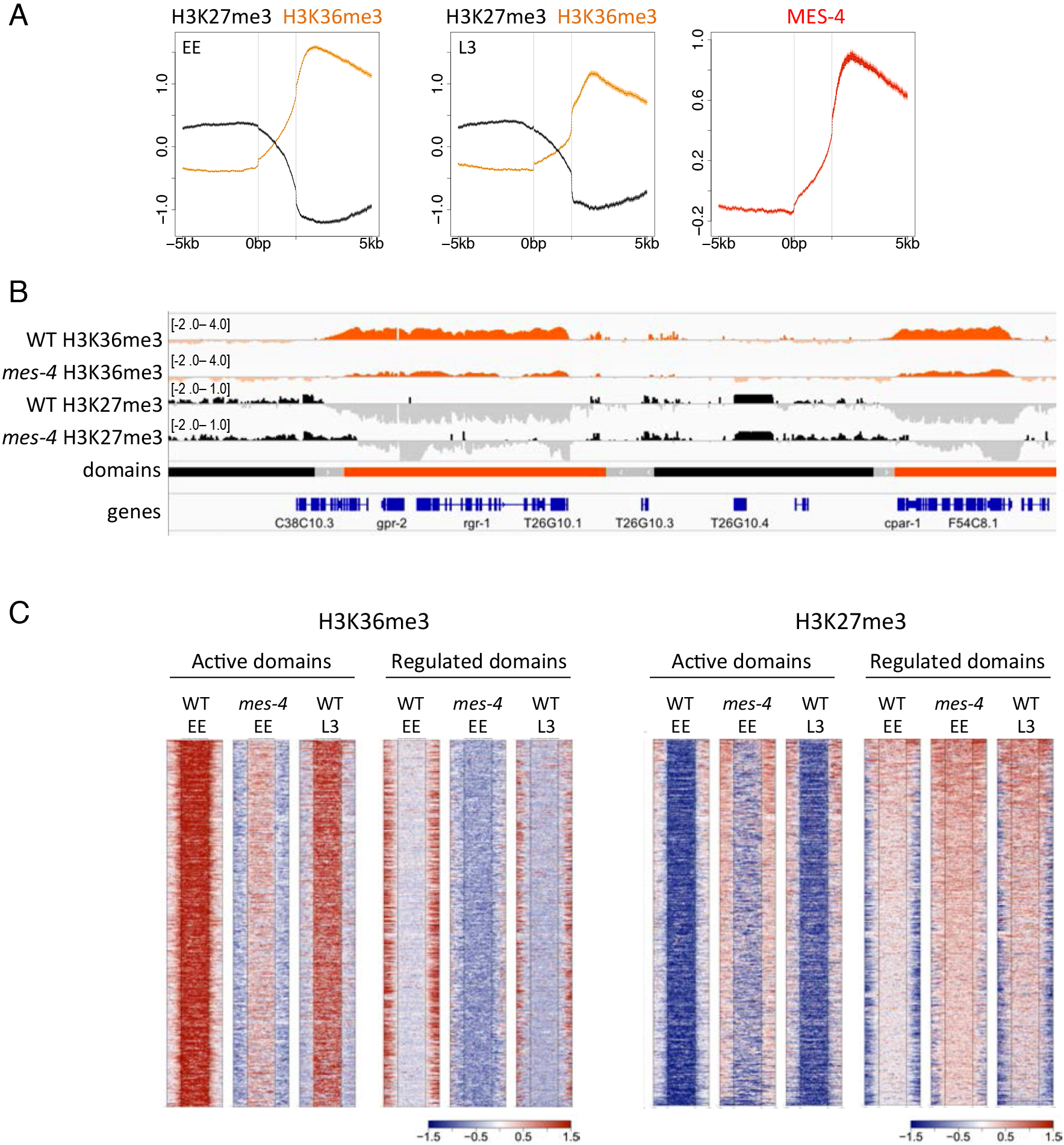
MES-4 regulates chromatin marking at domain edges. (A) Plots of H3K36me3 and H3K27me3 across regulated, border, and active domains in EE and L3, and of MES-4 in EE. Plots are centered at borders pseudoscaled at 2.5kb, and show 5 kb into regulated domains (left) and 5kb into active domains (right). Lines show mean signal, darker filled areas show standard error, and lighter filled areas are 95% confidence intervals. Grey vertical lines indicate edges of the border region. (B) IGV screenshot showing H3K36me3 and H3K27me3 tracks in wt and *mes-4* RNAi early embryos. (C) Heatmaps comparing H3K36me3 and H3K27me3 signals across EE active domains and EE regulated domains in wt and *mes-4* RNAi early embryos. Signals are centered on active domains or regulated domains (pseudoscaled at 5kb) as indicated, and plot 2.5kb into borders on either side.

The previous studies of *mes-4* focused on patterns of chromatin marking on individual germ line genes (19, 21). To investigate whether MES-4 is important for domain definition, we analysed patterns of H3K36me3 and H3K27me3 on domains in wild type and *mes-4* RNAi early embryos (using data from ref. 19). We found that H3K36me3 still marks active domains in *mes-4* RNAi embryos, but both the level and extent of H3K36me3 coverage over active domains is reduced (Figs. 3B,C). Complementing the reduction of H3K36me3 over active domains, we observed that H3K27me3 coverage at regulated domains is expanded (Figs. 3B, C). Because MES-4 is a germline H3K36 methyltransferase, these results suggest that chromatin regulation in the germ line contributes to the definition of active and regulated domains.

### Developmental remodeling of H3K27 and H3K36 methylation at domain borders

We next investigated whether patterns of H3K36 and H3K27 methylations in domains or borders were developmentally regulated. We observed H3K36me3 marking at borders is altered between EE and L3. In contrast to the sharp rise in H3K36me3 levels across EE borders, in L3 H3K36me3 levels are relatively low and constant in border regions (Fig. 4A). Although H3K27me3 patterns at borders are not obviously changed between EE and L3 (Fig. 4A), there is a striking remodeling of H3K27me1 patterns. In early embryos, borders have a strong peak of H3K27me1 enrichment, with lower levels in neighboring active and regulated domains (Fig. 4B). At the L3 stage, the H3K27me1 pattern is dramatically altered, with high levels at active domain edges and within active domains (Fig. 4B). The remodeling of H3K36 and H3K27 methylation patterns at border regions suggest that this regulation may play a role in domain definition.

**Figure 4.**
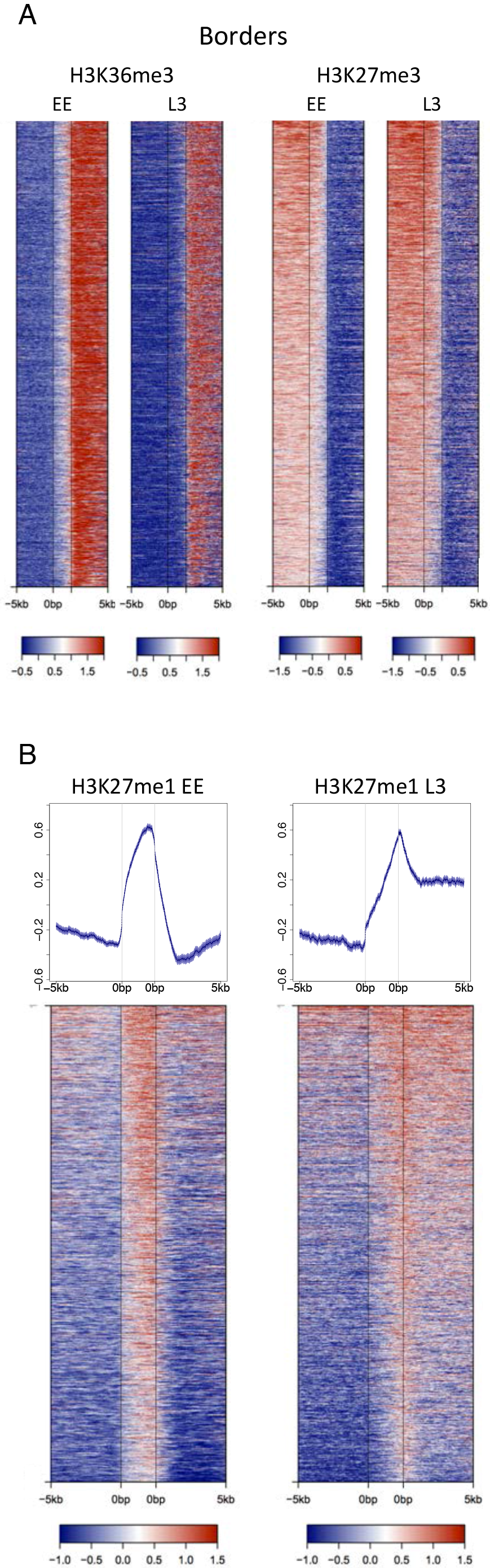
Remodelling of H3K36me3 and H3K27me1 marking from EE to L3. (A) Border H3K36me3 is reduced from EE to L3. Plots show heatmaps of H3K36me3 and H3K27me3 signals centered at borders pseudoscaled at 2.5kb, and show 5kb into regulated domains (left) and 5kb into active domains (right). (B) H3K27me1 at borders is remodeled from EE to L3. Upper plots average signals centered at borders pseudoscaled at 2.5kb, with 5 kb into regulated domains (left) and 5kb into active domains (right). Lines show mean signal, darker filled areas show standard error, and lighter filled areas are 95% confidence intervals. Grey vertical lines indicate edges of the border region. Lower panels show heatmaps of the same regions. Bottom plots show heatmaps of H3K27me1 signals centered at borders pseudoscaled at 2.5kb, and show 5kb into regulated regions (left) and 5kb into active regions (right). Heatmap rows are sorted by decreasing mean signal and show the same regions and order.

### Long intergenic regions and enrichment of transcription factor binding in borders

We observed that border regions were enriched for enhancer chromatin states and transcription factor binding sites, features of transcription regulatory regions. To further investigate the regulatory potential of border regions, we asked whether intergenic regions in borders have different properties than those in active or regulated domains. Comparing intergenic lengths in different regions, we found that those at borders are longer than those in active or regulated domains (Fig. 5). We also observed that intergenic regions in active domains are short compared to those in regulated domains (Fig. 5). Therefore, intergenic regions vary in length according to chromatin domain type across the genome. Borders, which mark the transitions between active and regulated chromatin domains, have the longest intergenic regions.

**Figure 5.**
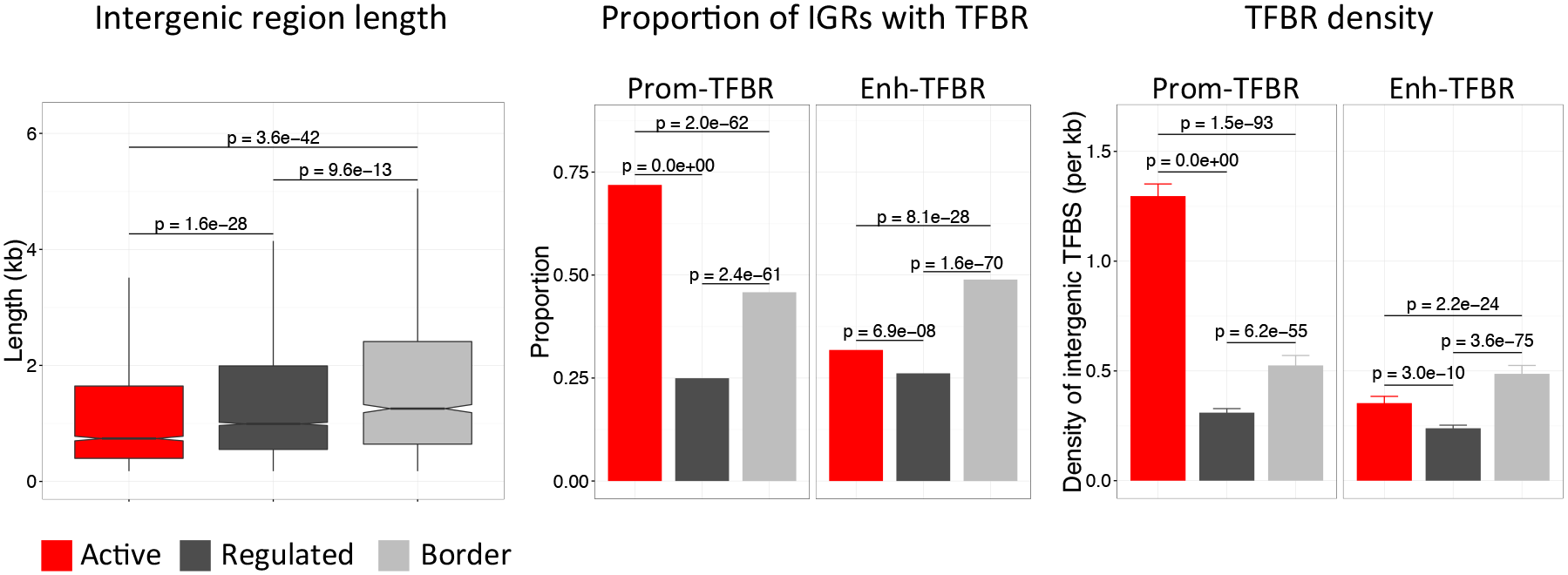
Intergenic regions at borders are long and enriched for enhancer TF binding sites. (A) Boxplots of intergenic region lengths in the specified regions; p-values show significance of distribution differences tested using the Mann-Whitney U test. (B) Proportion of intergenic regions overlapping at least one TFBR of the indicated type; p-values show differences in proportions tested using a Z-test. (C) Density of Prom-TFBR, and Enh-TFBR per kilobase of intergenic region; p-values show significance of distribution differences tested using the Mann-Whitney U test.

We next investigated the density of regulatory elements in intergenic regions in different domains. For this analysis, we separated 35,062 modENCODE transcription factor binding regions (TFBRs) (8, 9, 11) into two classes: those containing a promoter (Prom-TFBR; n=8388) and those not containing a promoter, which are likely enhancers (Enh-TFBR, n=26674). Intergenic regions within active domains more often have a Prom-TFBR and have a higher density of Prom-TFBRs than intergenic regions in borders or regulated domains (Fig. 5). In contrast, border intergenic regions more often contain an Enh-TFBR and have a higher density of Enh-TFBRs than those in active or regulated domains (Fig. 5). Therefore, intergenic regions in borders are longer and more enriched for Enh-TFBRs compared to those in domains. The location of extended transcriptional regulatory regions between active and regulated chromatin suggests that transcriptional activity may play a role in separating chromatin domains.

## DISCUSSION

Here we derived chromatin state maps for two developmental stages of *C. elegans* and used the resulting chromatin states to investigate chromatin domain organization. We show that *C. elegans* autosomes are subdivided into extended chromatin domains of differing activity separated by border regions containing long, regulatory element rich, intergenic regions. Chromatin domain positions are remarkably consistent between early embryos and L3 larvae, despite the two stages being comprised of nearly non-overlapping cell types. Therefore, chromatin domain organization appears to be a basal property of the genome.

Figure 6 shows a simple model to explain our observations. The two types of chromatin domains contain different types of genes and are subject to different modifications. “Active” domains contain broadly expressed genes and “regulated” domains contain genes that are developmentally regulated or that have low or undetectable expression. Genes in active domains are expressed in the germ line and there are subject to H3K36me3 marking by MES-4 (21–23). This modification is inherited and maintained in early embryos by maternally provided MES-4. When gene expression is zygotically activated in the embryo, these genes would be marked by the transcription coupled H3K36 histone methyltransferase MET-1 (20–22), preserving this pattern somatically. Regulated genes and those with no/low detectable expression lie in regulated domains that are marked by H3K27me3. These genes show low or no H3K36me3 modification. Although the profiling in mixed tissues done here may have limited the ability to detect tissue specific H3K36me3, the results suggest that regulated genes acquire no or only low levels of H3K36me3 when they are transcribed. In support of this possibility, a recent study showed that regulated genes are not marked by H3K36me3 when they are expressed (24). We propose that H3K36me3 marking may be specifically relevant to genes with stable expression across development, possibly aiding the stability of their expression. Consistent with this idea, it was recently demonstrated that H3K36me3 marking plays a role in gene expression stability during aging in *C. elegans* (25). H3K36me3 could also play a role in the preservation of chromatin domain structure.

**Figure 6.**
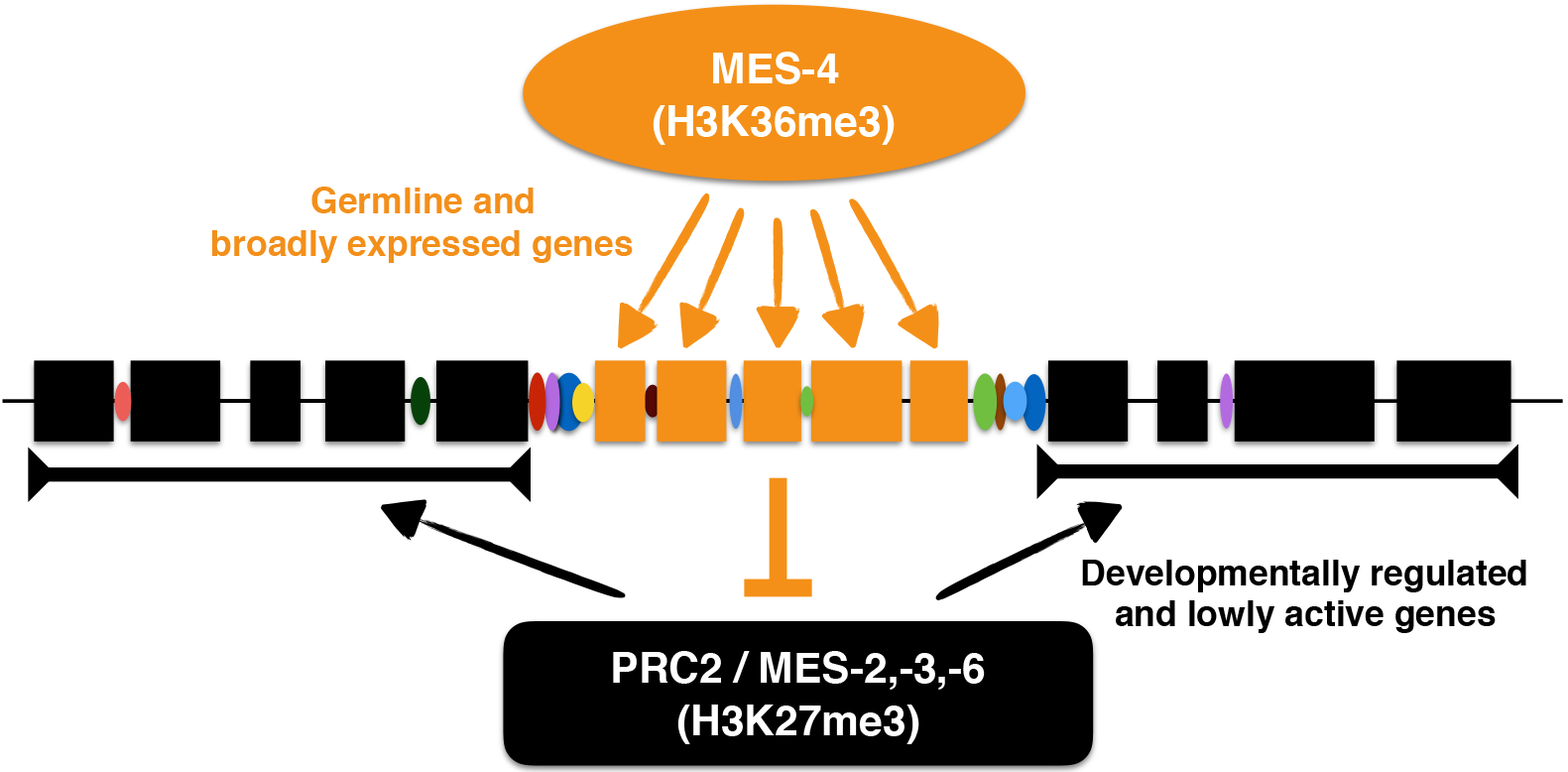
Model for regulation of chromatin domains by MES-4, PRC2, and transcription regulatory regions. The genome is subdivided into domains of germline and broadly expressed genes (orange boxes), and domains of regulated and lowly expressed genes (black boxes). MES-4 marks germline expressed genes with H3K36me3, which inhibits deposition of H3K27me3 by PRC2 (19, 21), leading to demarcation of chromatin domains. Borders separating domains contain long intergenic regions enriched for transcription factor binding (colored ovals); these regulatory regions may play a role in domain separation.

The Polycomb repressive complex PRC2 generates H3K27me3, H3K27me2, and H3K27me1, although it is not well understood how the different levels of modification are regulated (23, 26–30). A large body of work has shown that PRC2 functions to maintain and propagate transcriptional repression, but it can also be permissive for transcriptional activation (31–33). Similar to the patterns analyzed here for *C. elegans*, broad domains of H3K27me3 or H3K27me2 laid down by PRC2 and that anticorrelate with high gene activity or H3K36me3 have been observed in other organisms (19, 28, 34–37). This anticorrelation is consistent with the inhibition of PRC2 activity by H3K36me3 and other histone modifications associated with gene activity (26, 27, 29, 30, 38, 39). These patterns suggest that PRC2, together with features of active chromatin such as H3K36me3, and interactions between them, may play a conserved role in the formation of domains of differing chromatin activity.

Although the experiments performed here on whole animals (early embryos and larvae) captured the high similarity of chromatin domains in development, the mixed tissues precluded our ability to study how chromatin domains might be regulated in individual cell types. Performing similar studies using purified cell types would be needed to investigate this question.

The features of the border regions between active and regulated domains suggest a role for transcription regulation in separating domains. We observed that intergenic regions at borders are generally longer than those in active or regulated domains and they are more enriched for TF binding sites distal from promoters, which are likely to define enhancers or other regulatory elements such as insulators. These properties of borders combined with their location between chromatin domains of different activity suggest that transcription regulatory regions may be involved in domain separation. For example, the binding of factors to borders might generate a blocking structure or a platform for interactions. It is also possible that transcriptional activity or the generation of chromatin accessibility is important. Border regions show high chromatin accessibility, as do functional boundaries even when neighboring genes are not active (40–44). Mechanisms operating at border regions could also act in conjunction with those actively specifying domains.

An important future goal will be to understand the relationship between chromatin domains, which reflect chromatin activity, and three-dimensional structure. Using techniques to measure chromatin interaction frequencies (“C” methods), it has been shown that chromatin has different levels of three-dimensional organization within the nucleus, from broad chromosome territories to smaller scaled topologically associated domains or TADs (43, 45–48). Topological domain mapping gives information about overall genomic structure, but not underlying genomic activity, whereas chromatin state mapping provides information on chromatin composition, but not on physical interactions. A Hi-C chromatin interaction map for *C. elegans* was recently published, defining 10-20 large domains per chromosome each containing an average of ~200 genes (48). This is much higher than the 7-12 genes in human, mouse and Drosophila TADs (43, 45–47), suggesting that these *C. elegans* Hi-C domains may be functionally different. Because the chromatin domains defined here contain an average of five to nine genes (median of four), the large *C. elegans* Hi-C domains would each harbor many chromatin domains. Unfortunately, the resolution of the *C. elegans* Hi-C domain boundaries (10 kb) is currently not sufficient for a comparison with the border locations defined here (48). Future higher resolution chromatin interaction studies will be needed to determine how the chromatin domains studied here relate to three-dimensional structure.

In summary, our results point to roles for germ line chromatin marking and transcription regulatory regions at chromatin domain borders in organizing the genome into functional domains and provide a framework for studies of chromatin structure and function in *C. elegans*. The future identification and functional analyses of sequences and factors that control chromatin structure will allow a better understanding of the mechanisms and functions of genome domain organization.

## Materials and Methods

### Datasets and processing

The datasets used for generating the chromatin state maps were early embryo and third larval stage ChIP-chip or ChIP-seq histone and histone modification data (7, 10). Data are available from Gene expression omnibus and http://data.modencode.org/ (Table S2 gives GEO accession numbers). For ChIP-chip data, log ratios of experiment signal over input signal were normalised, z-scored and then averaged over replicates. Probes assigned to repeat regions were omitted. ChIP-seq data were processed using BEADS (49) at 1bp resolution and averaged for the matching 50 bases of the ChIP-chip probes then logged and z-scored. The data was then corrected for outliers by considering a moving window of 9 probes: the central value was replaced by the average of the adjacent values, if (| x- m |) / s > 3, where x is the central value and m and s are the mean and sample variance of the remaining eight values. The data was reduced to the set of probes for which data was available for all 17 marks.

Throughout, genome coordinates used were WS220. Other data used: *C. elegans* WS224 gene positions lifted over to WS220 coordinates (www.wormbase.org); operon annotations from Wormbase WS220; definition of arm and center chromosomal regions (14); categories of non-coding genes from WS220; odorant receptors (hand curated list from C. Bargmann based on (50)); tissue specific genes (n=748), genes core enriched in neurons, intestine, hypodermis, body wall muscle, or coelomocytes from (51); repeats from Dfam2.0 (52); ubiquitous genes (n=2575) (21); promoter and enhancer annotations (53); RNA Polymerase II ((11), EE: GEO GSE25788, L3: GEO GSE25792); MES-4 ((19), GEO GSE38180); H3K36me3 and H3K27me3 in wt and *mes-4* RNAi EE ((19), GEO GSE38180 and GSE38159); stage specific RNA-seq data (54). Silent genes were defined as those with <0.005 dcpm in EE, LE, L1, L2, L3, L4, and YA hermaphrodite RNA-seq from (54) (n=1921). Maternally expressed genes were defined using Cel-seq data profiling the AB and P1 blastomeres of two cell stage embryos (55). RNA in AB and P1 blastomeres is maternally contributed (and therefore germline expressed) since these blastomeres have negligible gene expression. Genes with mean rpkm>0 in both AB and P1 were classified as maternal (n=7980). Genes with broad or regulated expression were defined based on gene expression variability scores, which are the coefficients of variation (cv) in gene expression (ratio of the standard deviation and mean expression) across 35 samples of different stages and cell types; cv values are from (54). A low cv value indicates a gene has similar expression across all samples (broad expression) whereas a high cv indicates a gene has high variation in expression across samples (regulated expression). As genes with very low expression often have high cv values, we only considered those with moderate to high expression in at least one developmental stage (dcpm >0.2, n=13739). Genes in the bottom third of these cv values were defined as having broad expression and genes in the top third of cv values were defined as having regulated expression. Metagene plots and heatmaps were generated using Seqplots (http://przemol.github.io/seqplots/). The IGV Integrative Genome Viewer was used to visualize data (56, 57).

### Generating the chromatin state models

We chose 20 states as a practical compromise between capturing the complexity of biological features and ease of interpretation; models with a larger number of states contained states that were superficially similar to each other. The states were found using a standard Hidden Markov Model for multivariate gaussians (each state having parameters for mean and covariance of the marks), with version 1.0.4 of RHMM (http://cran.r-project.org/) (58), with runtime parameters: nStates=20, dis = "NORMAL", control=list(verbose=2,init="KMEANS", iter=500). As whole genome state maps separated into autosomal and X chromosome-specific states, we generated autosomal and chromosome X chromatin state maps for each stage: EE autosomes, EE Chr X, L3 autosomes, and L3 Chr X. For each, forty replicate chromatin state HMMs were generated from different random number seeds. To assess the consistency of the 40 replicates, we first matched states between every pair of replicates so that they are comparable. The Jaccard Index (ratio of the length of intersection between two regions to the union of genomic coverage of the two regions) was calculated between every state in one replicate and every state in the other, forming a 20x20 similarity matrix from which the pair of states with highest similarity was matched iteratively until all states were matched. Then an overall Jaccard Index was calculated between every pair of replicates either from the same stage (e.g, between matched states in the 40 EE autosome replicates) or between stages (e.g., EE autosome to L3 autosome). Within a stage, the replicate 20 state models were very similar to each other (Figure S2A); one from each set was chosen for analysis (EE autosome, EE Chr X, L3 autosome, L3 Chr X). Chromatin states were matched between the chosen EE autosome and L3 autosome map and between the EE Chr X and L3 Chr X maps by maximizing the Jaccard Index. Figs. S2B,C shows the similarity between individual EE and L3 autosomal chromatin states in the chromatin state maps analysed in this paper.

As the chip probes are 50 bases long and 147 bases wrap around one nucleosome, chromatin states of one or two probes were considered too short to be biologically meaningful and so reassigned to adjacent states. Segments of exactly one probe were assigned alternately to the left or right state: segments of exactly two probes were split, the first probe assigned to the left state, the second to the right. Where a continuous instance of a state was defined, for example for statistics on length of states, a run of consecutive probes of the same state including gaps between probes up to a gap of 500 bases was used. Probes on either side of larger gaps were treated as being in different individual states. Dataset S1 gives coordinates of the 20 states in each of the four maps. In feature charts, the cells show fold enrichment on a log2 scale. Cells were colored grey if there were too few data points for statistical confidence: if at 95% confidence there were too few data points to be sure that the estimate was within half a unit on the log2 scale of the real value.

### Definition of highly active and lowly active domains

Autosomal domains were defined from the chromatin states as follows: States 1-5 (the most strongly associated with the highest quintile of gene expression) were defined as active, states 16-20 (the most strongly associated with the lowest quintile of gene expression) were defined as inactive, and states 6-15 were defined as neutral. Regions containing active states and the neutral states among them without interruption by states 16-20 were defined as highly active domains (later renamed “active domains”), ending with an active state. Regions containing inactive states and the neutral states among them without interruption by states 1-5 were defined as lowly active domains (later renamed “regulated domains”), ending with an inactive state. Regions between active and regulated domains were defined as borders. Single active states between two inactive states and single inactive states between two active states were considered neutral to prevent singleton states breaking up domains; this occurred in only a small number (~10%) of domains. Domains with less than 50% of their length covered by states were removed. Dataset S2 gives coordinates for regulated, border, and active regions in EE and L3. Tables S4 gives the number of domains of each type, lengths, and number of genes per domain, along with expected numbers based on the simulation described below. To count the number of genes per domain, protein coding genes were assigned to the domain that overlaps its midpoint. The expected number of genes per domain was obtained by permutation, in which domains were called from randomly shuffled chromatin states (Table S1). To control for the positive effect of genic organization on domain length (e.g, association of promoter, transcription elongation, and gene end states, and operons), states associated with genes were shuffled as units and operon genes were shuffled together and considered single genes. The permutation was repeated 100 times.

### Intergenic region length and transcription factor binding analyses

Intergenic regions were defined as regions on autosomes between annotated protein coding genes (WS220), excluding those where genes overlap, and those separating genes in operons (n=13,705). To avoid biases caused by outliers, we excluded the top and bottom 10% of intergenic region lengths. Intergenic regions were assigned to the domain which had the largest value of reciprocal overlap as defined as (length_overlap)^2^ / (length_domain x length_IGR). TF binding regions (n=35,062) were merged modENCODE TF peak calls for 90 *C. elegans* factors, processed as Supplemental File S1 in (59); data are from (8, 9, 11). Promoter annotations were protein coding TSSs from (53) and (60). For genes with no TSS annotation in either set Wormbase TSSs were used. TF binding regions were annotated as promoters if they overlapped a TSS annotation (Prom-TFBR, n=8388). TF binding regions that did not overlap the TSS set were annotated as likely enhancers (Enh-TFBR, n=26674). TF binding regions were assigned to intergenic regions on the basis of simple overlap. Differences in the distributions of intergenic lengths and TF binding site densities in different chromosome domains were tested using the Mann-Whitney U test. We tested if the proportions of intergenic regions hosting TFBRs are the same in different chromosome domains using a Z-test.

## Acknowledgments

We thank R. Chen, T. Gaarenstroom, C. Gal, and J. Janes for helpful comments on the manuscript and P. Kharchenko for advice in the early stages of the project. We are grateful to C. Bargmann, J. Ho, and B. Alver, and L. Hillier for providing processed or curated data. NH, PS, MC, and JA were supported by a Wellcome Trust Senior Research Fellowship to JA (054523 or 101863). KJE and TAD were supported by a Wellcome Trust Research Career Development Fellowship to TAD (083563w). JA and TAD also acknowledge support by core funding from the Wellcome Trust (092096) and Cancer Research UK (C6946/A14492).

---

**Figure S1.**
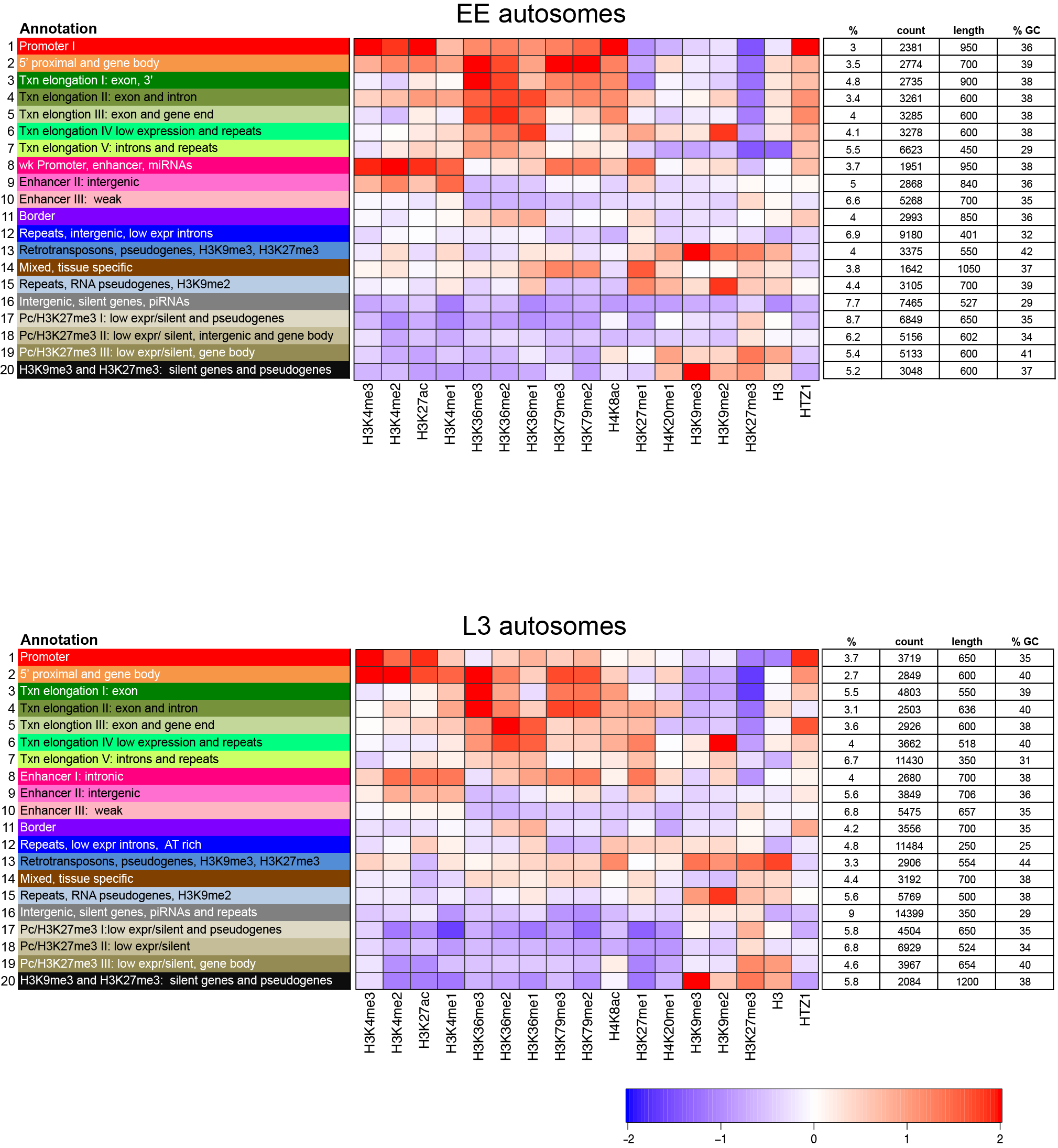
Chromatin state map keys and annotation for EE autosomes, L3 autosomes, EE chromosome X and L3 chromosome X. Left panel gives state numbers and annotations, middle panel shows relative enrichment or underenrichment of the indicated histones or histone modifications in each state, right panel gives for each state its % genomic coverage, number of instances, median length, and % GC. The scale bar shows the average z-score of the mark.

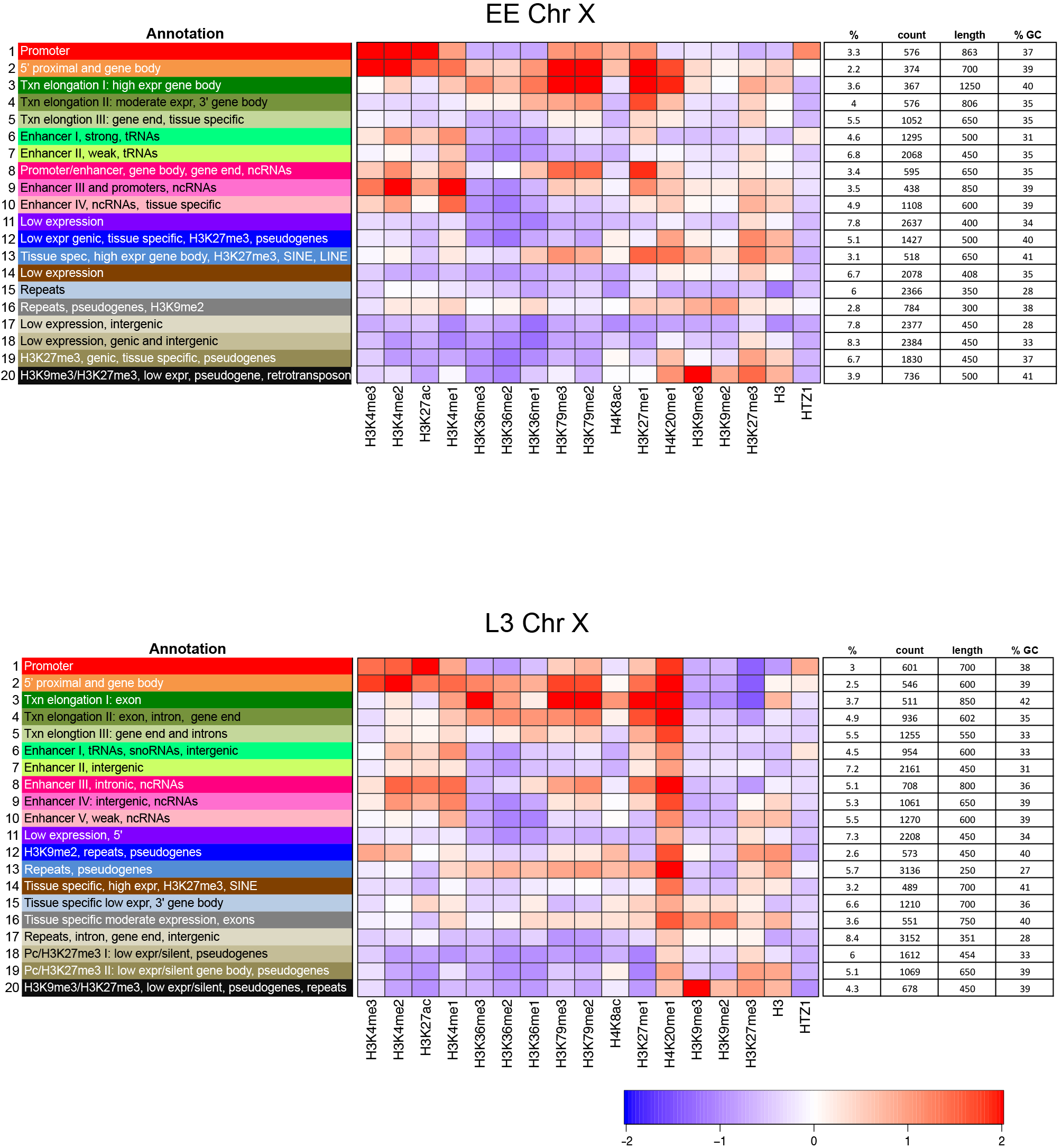

**Figure S2.**
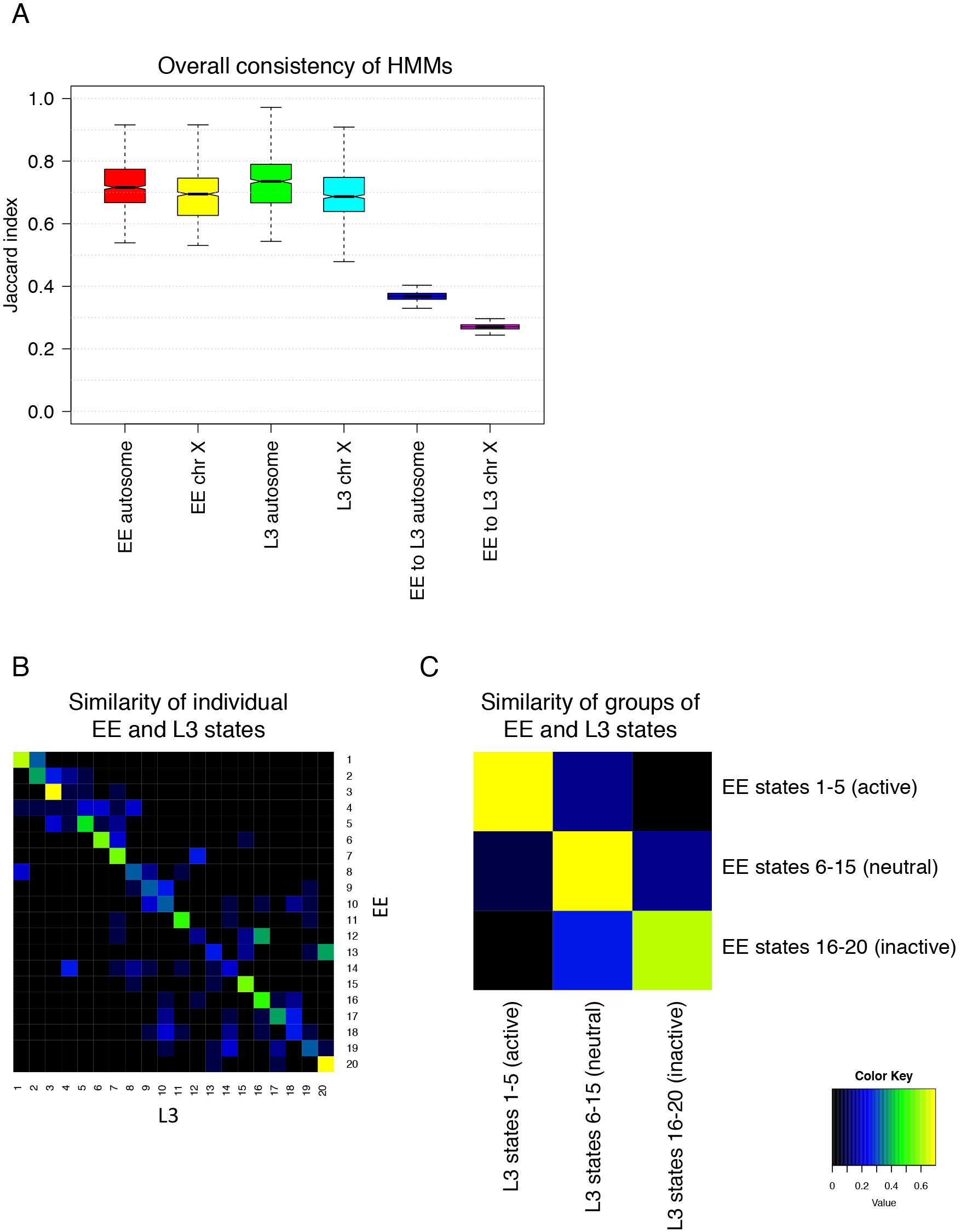
Consistency of chromatin state Hidden markov models (HMMs). (A) Left four boxplots show the distribution of Jaccard indices for 40 replicate runs of EE autosome, L3 autosome, EE chr X, and L3 chr X maps. Right two boxplots show distribution of Jaccard indicies between EE and L3 autosome replicates and between EE and L3 chr X replicates. (B) Comparison of the individual EE and L3 chromosome states in the maps analysed in this paper. (C) Comparison of EE and L3 chromatin states defined as active (states 1-5), neutral (states 6-15), and inactive (states 16-20). In general, states and state types are highly similar in EE and L3.

**Figure S3.**
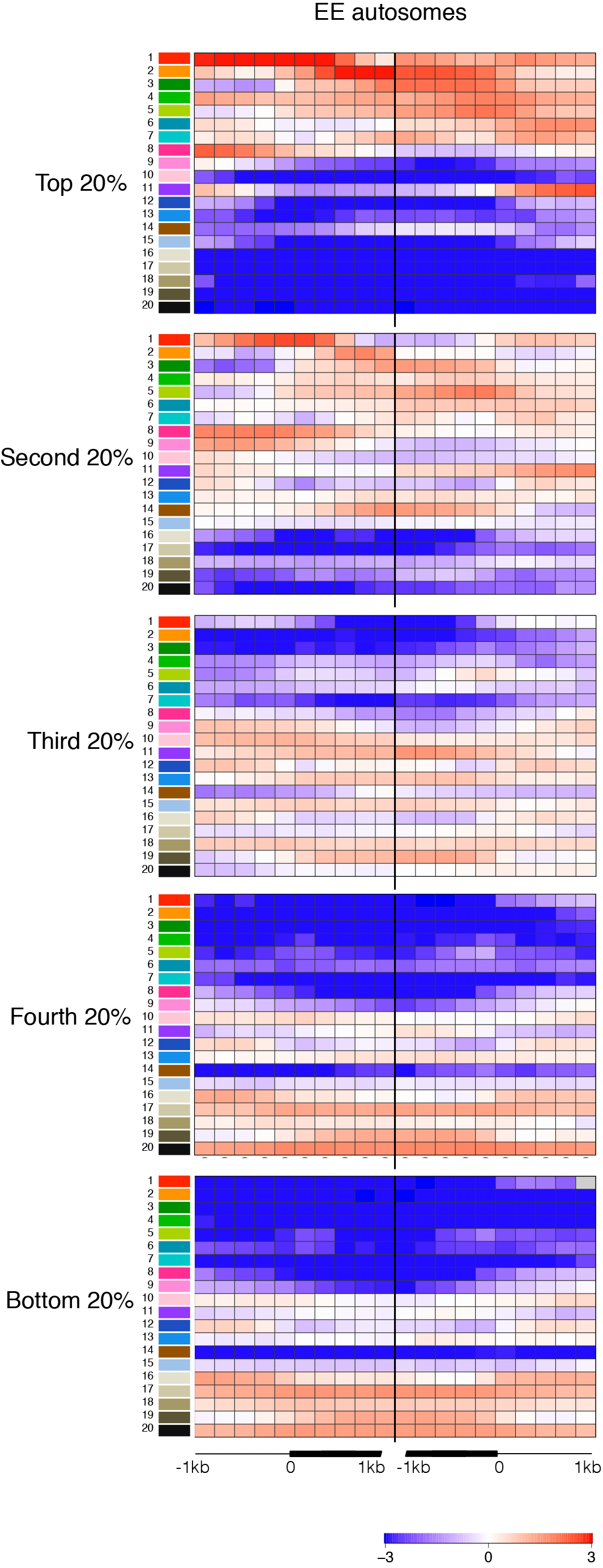
Chromatin state enrichments of genes across each quintile of gene expression for EE autosomes, L3 autosomes, EE chromosome X and L3 chromosome X. Enrichments are shown for 200bp windows spanning 2kb regions centered at Wormbase gene starts (left of black line) or ends (right of black line), as indicated by cartoon at the bottom. The scale bar shows log2 fold enrichment or underenrichment.

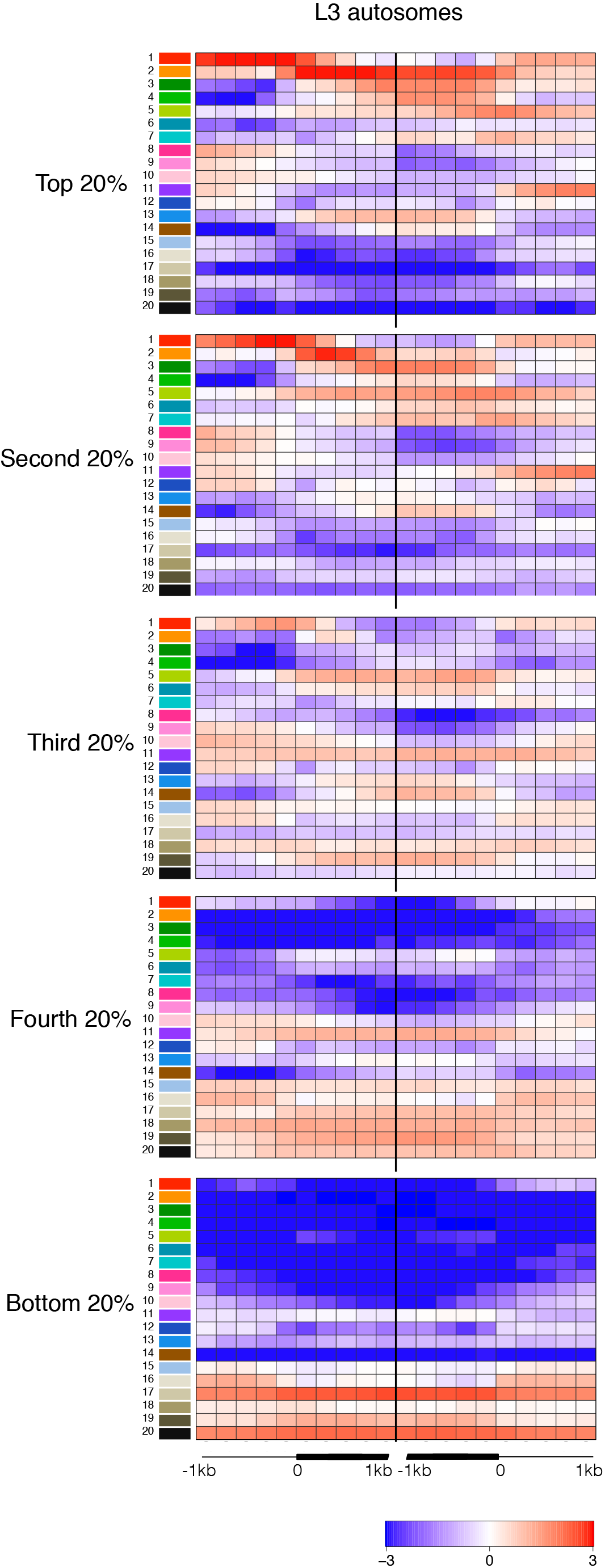

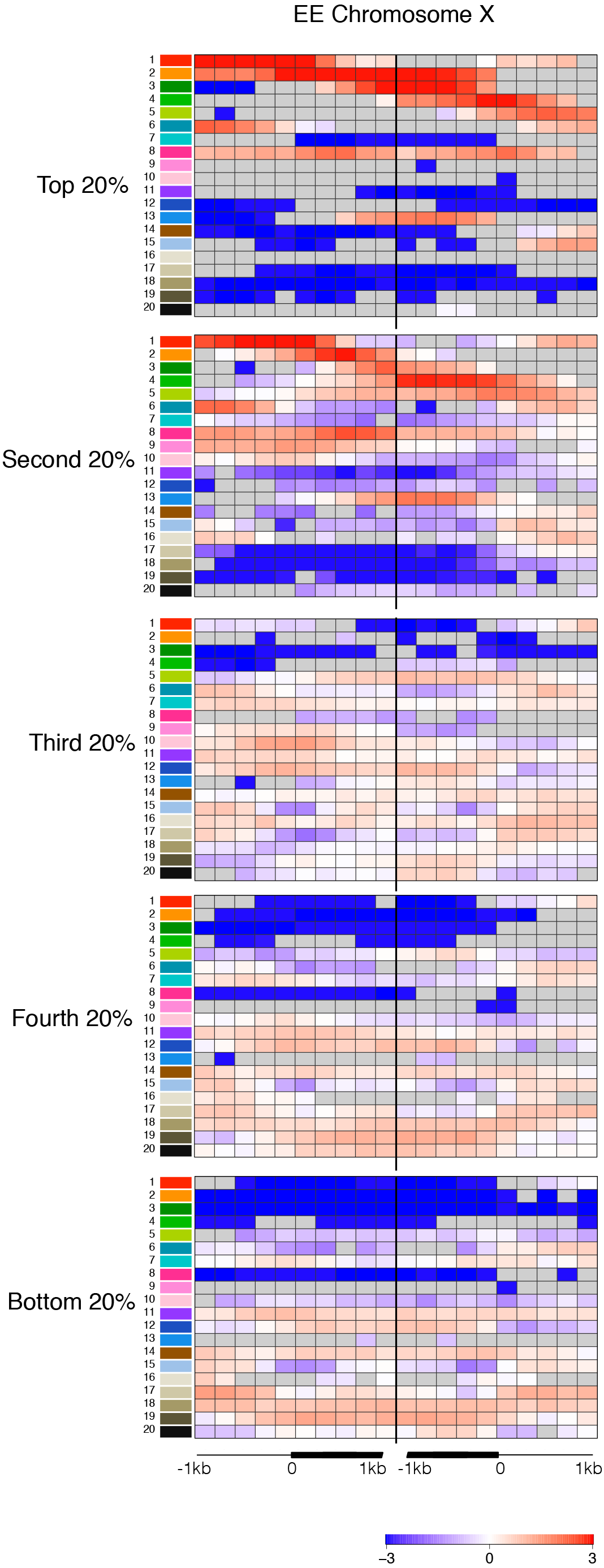

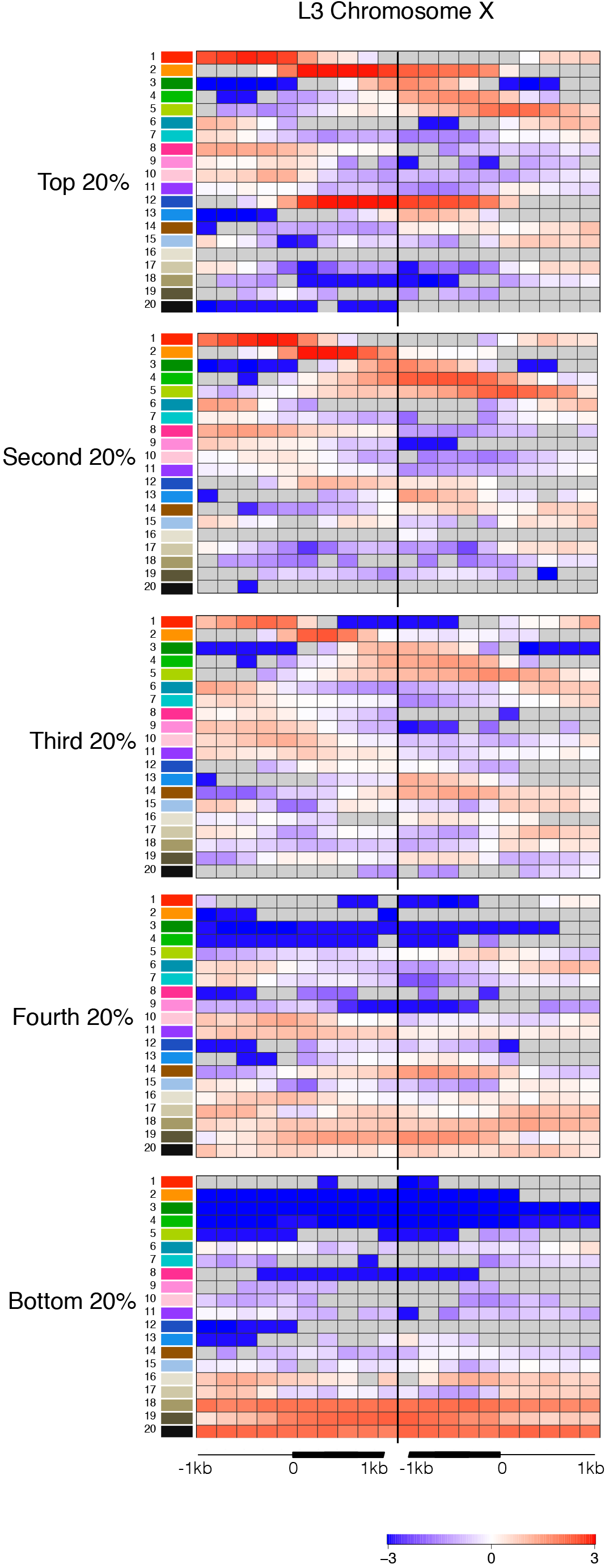

**Figure S4.**
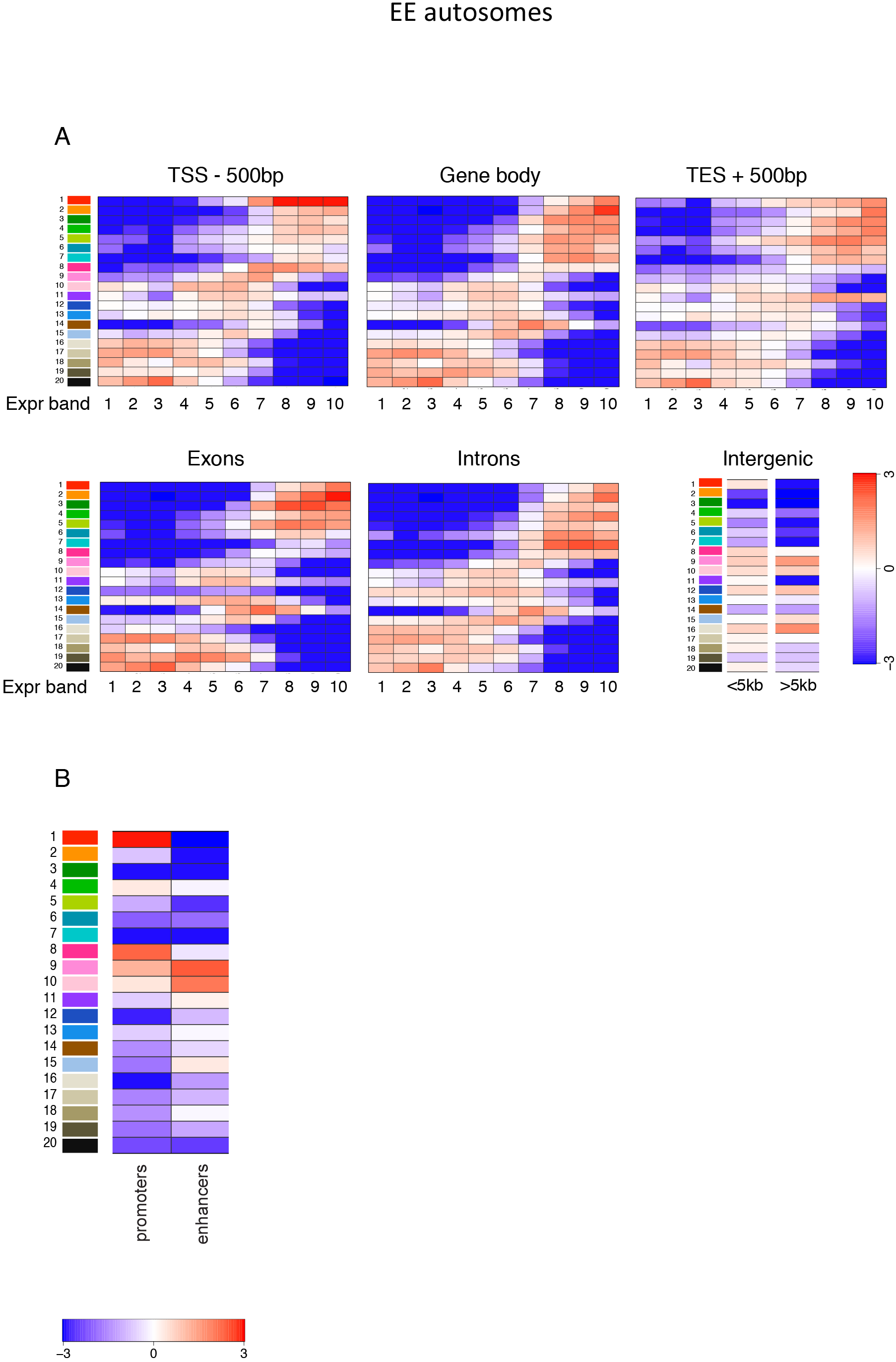
Chromatin state enrichments on different genomic features and features in EE autosomes. (A) State enrichments in different genomic features. For each feature, enrichments are shown separately for genes in each decile of expression (expression band 1=lowest). (B) State enrichments in promoters and enhancers. Promoter and enhancer definitions are from (53). (C) State enrichments in coding and different classes of non-coding RNAs. Class definitions are from Wormbase. (D) State enrichments on repeat elements. Repeat charts show state enrichments on all repeat, named classes, and a selection of individual repeats; repeat annotation is from Dfam2.0 (52). (E) State enrichments on groups of genes with low H3K27me3/low gene expression, low H3K27me3/high gene expression, high H3K27me3/ high gene expression, or high H3K27me3/low gene expression, or tissue specific genes. (F) Distribution of states on chromosomes, chromosome regions, odorant receptors, and pseudogenes. Cells are colored grey if there were too few data points for statistical confidence. Scale bars show log2 fold enrichment or underenrichment.

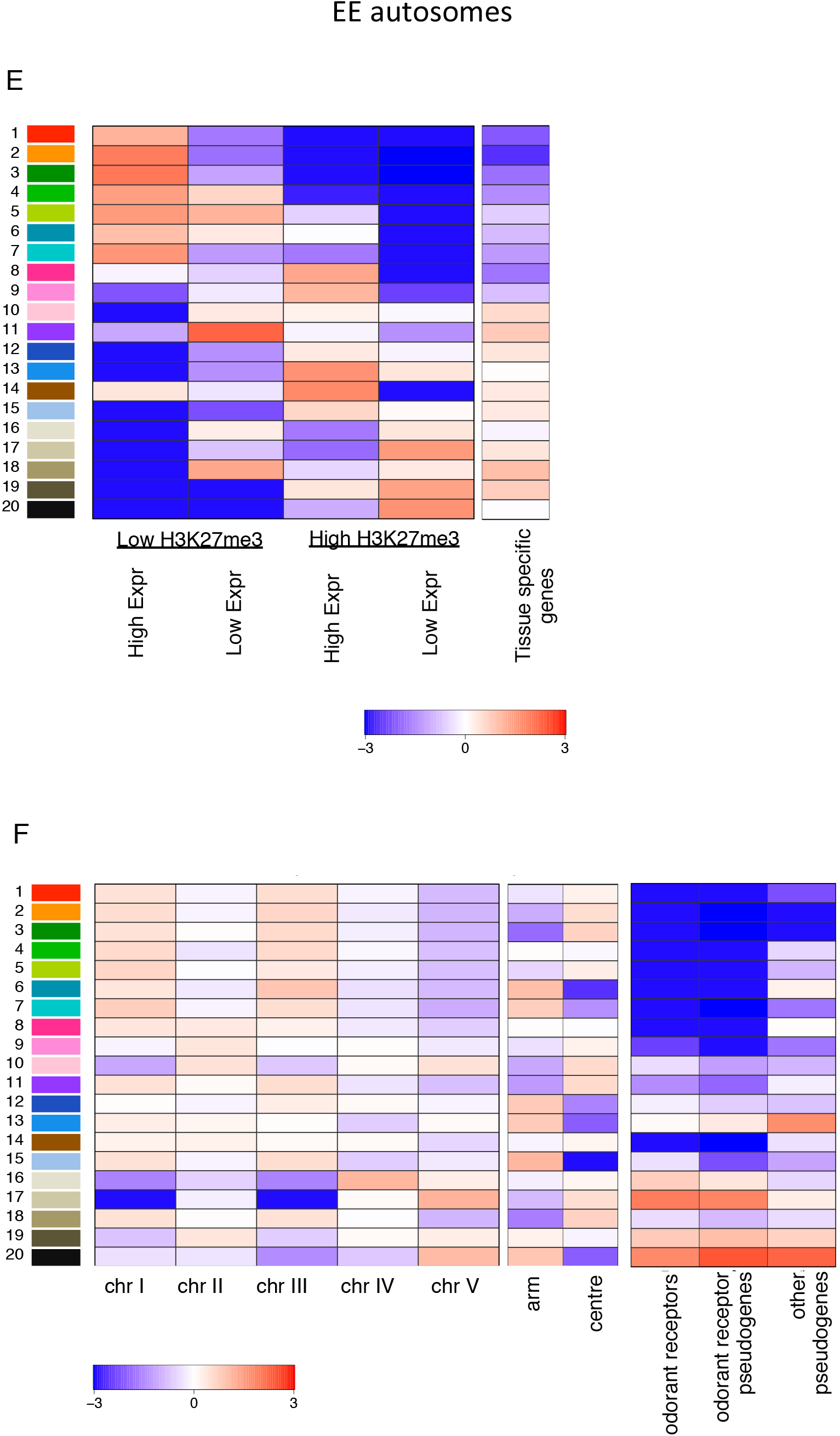

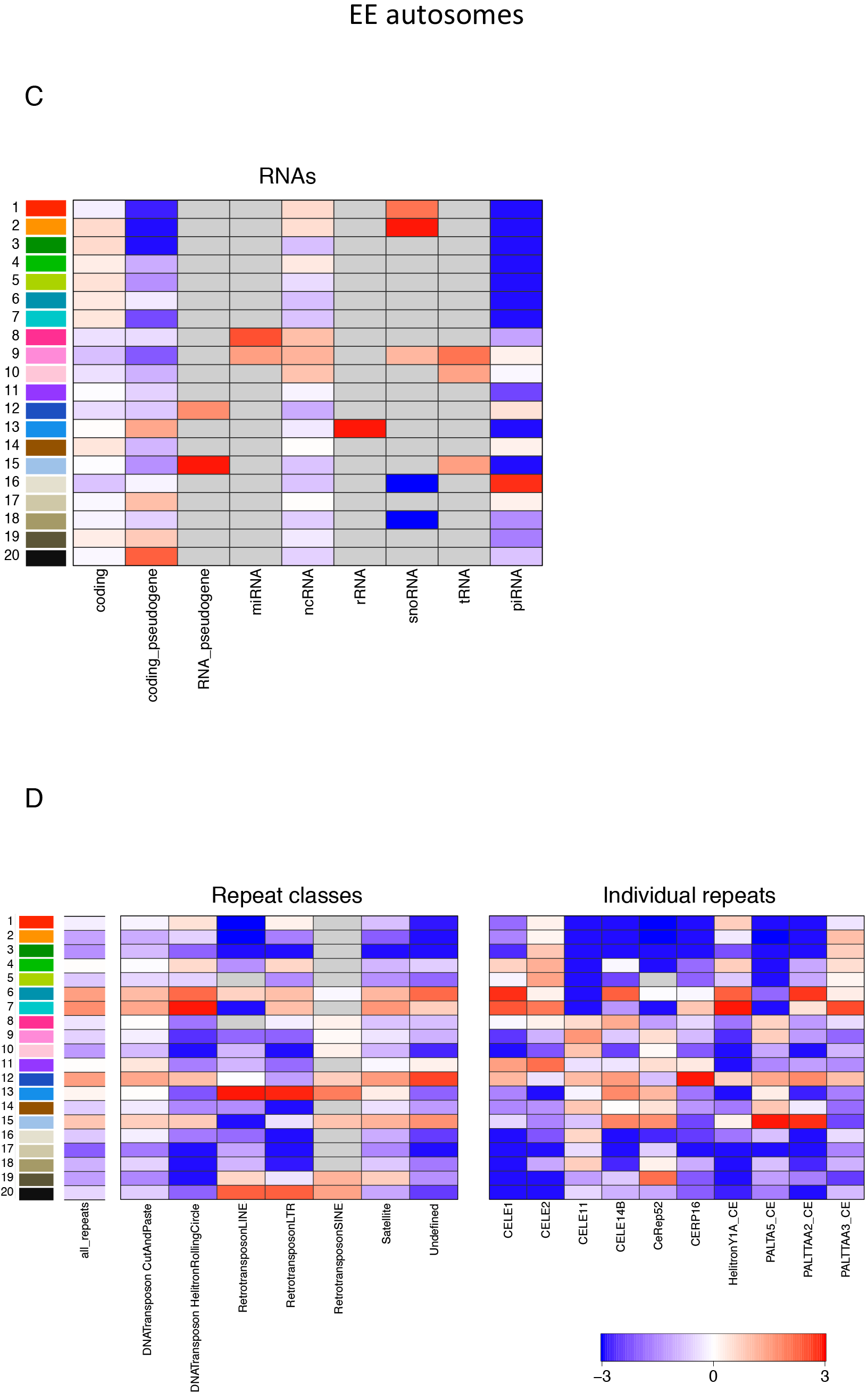

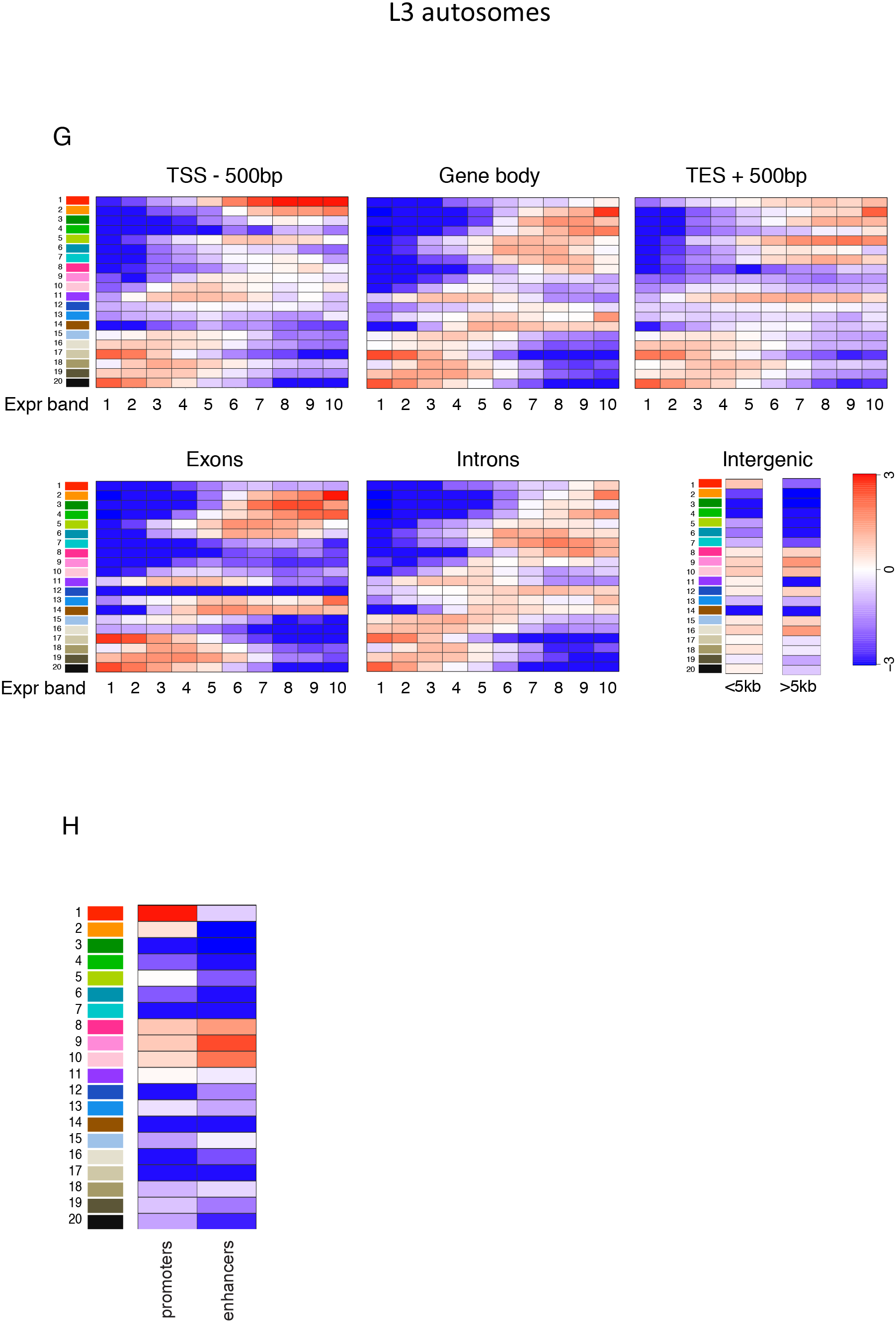

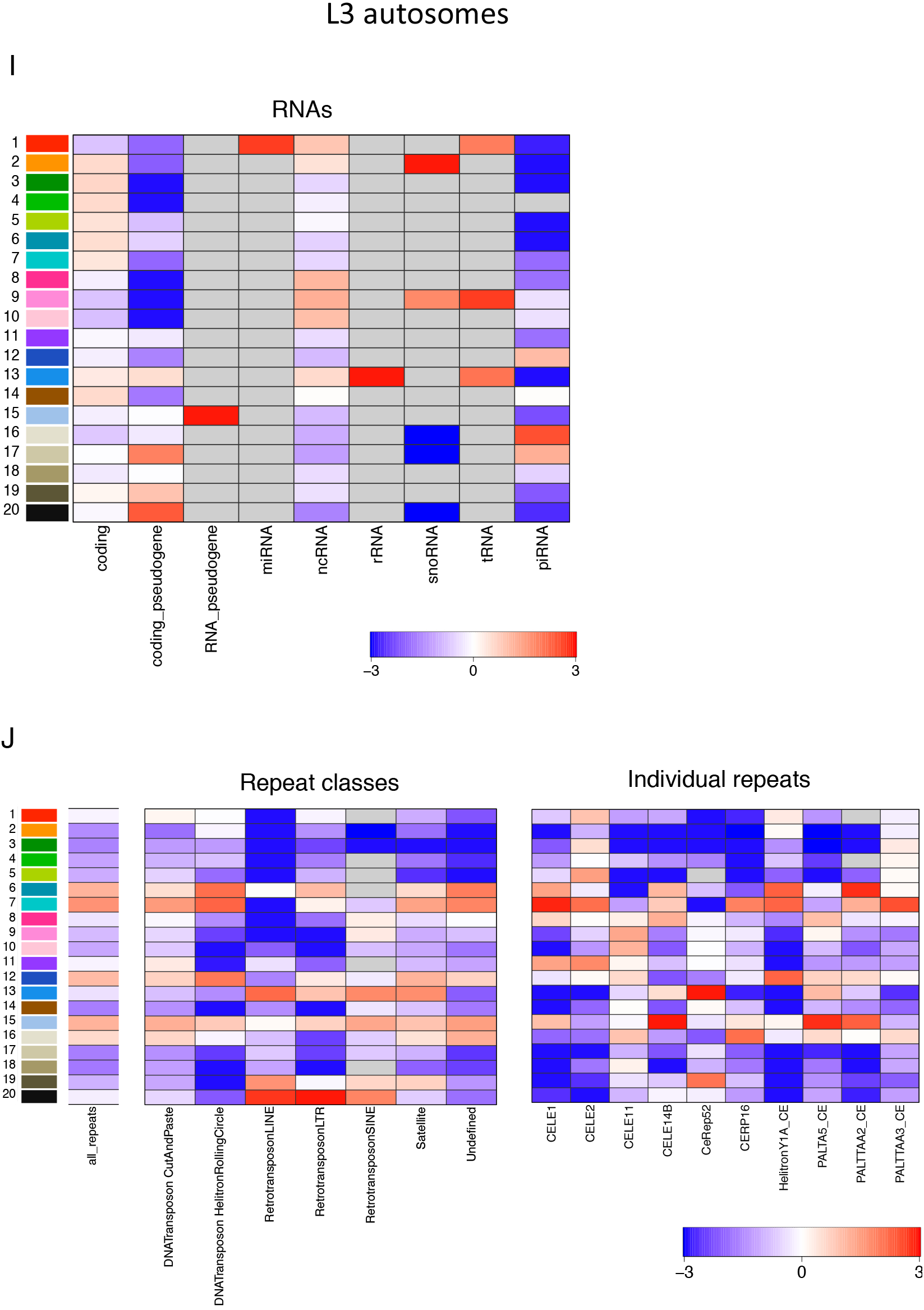

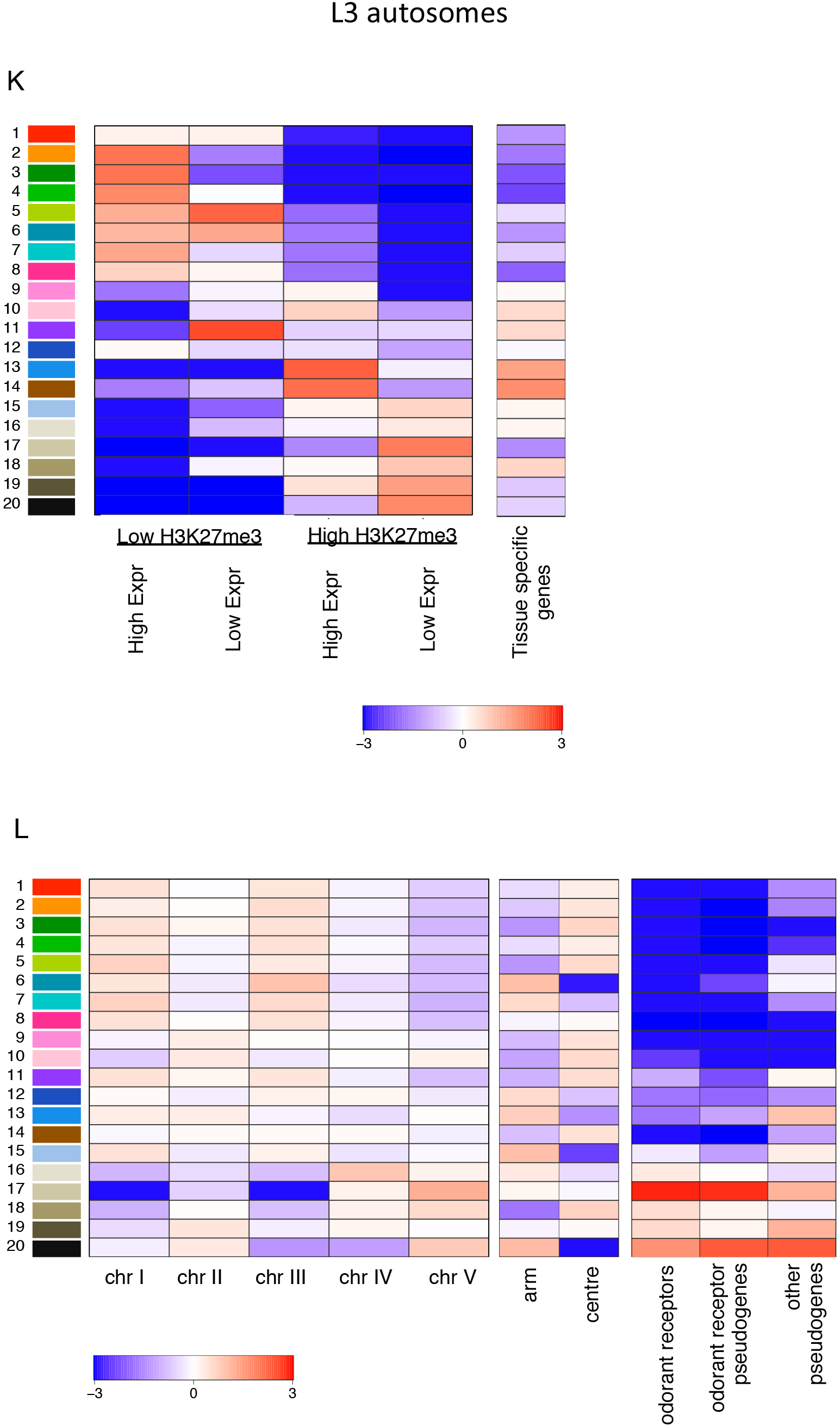

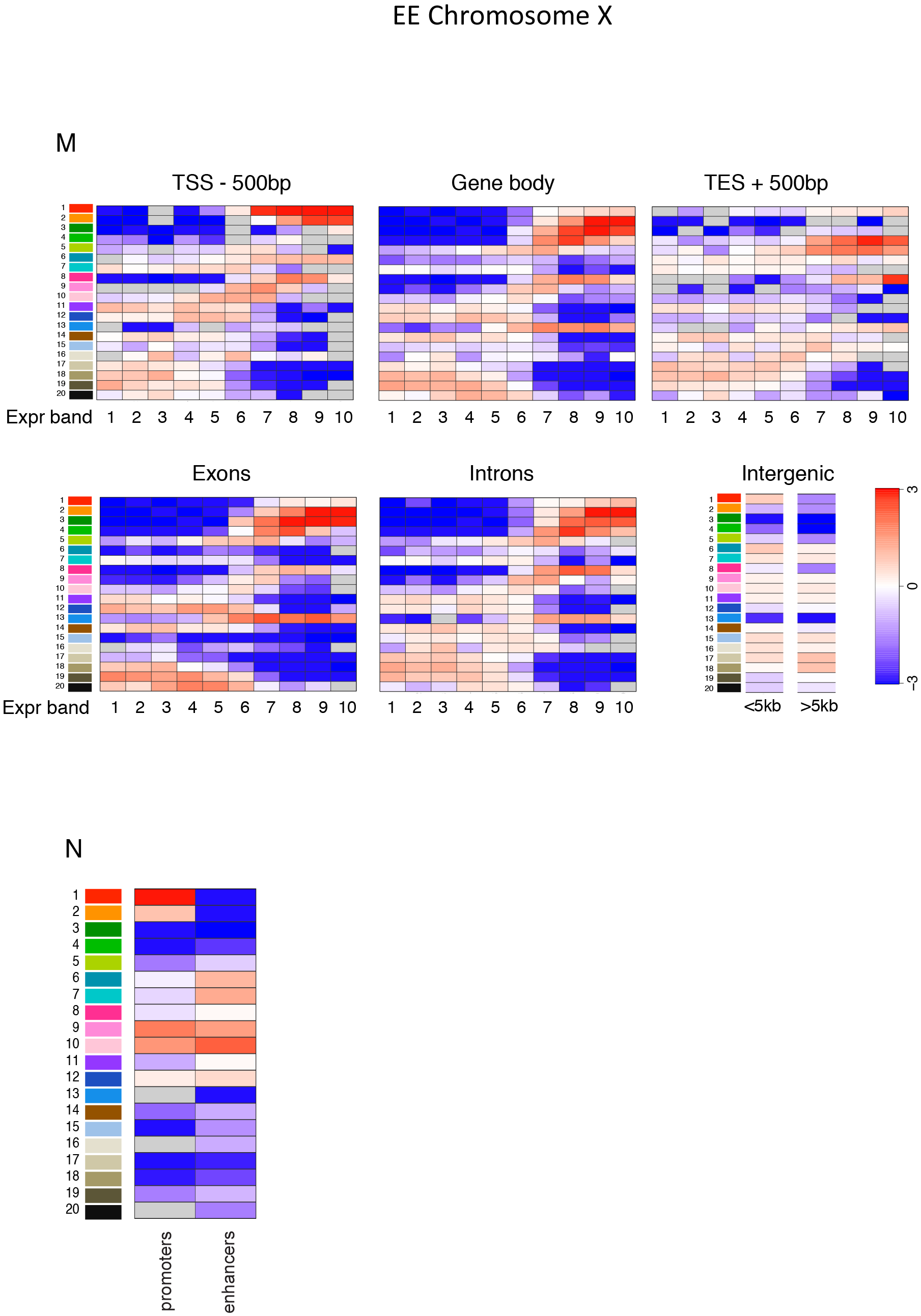

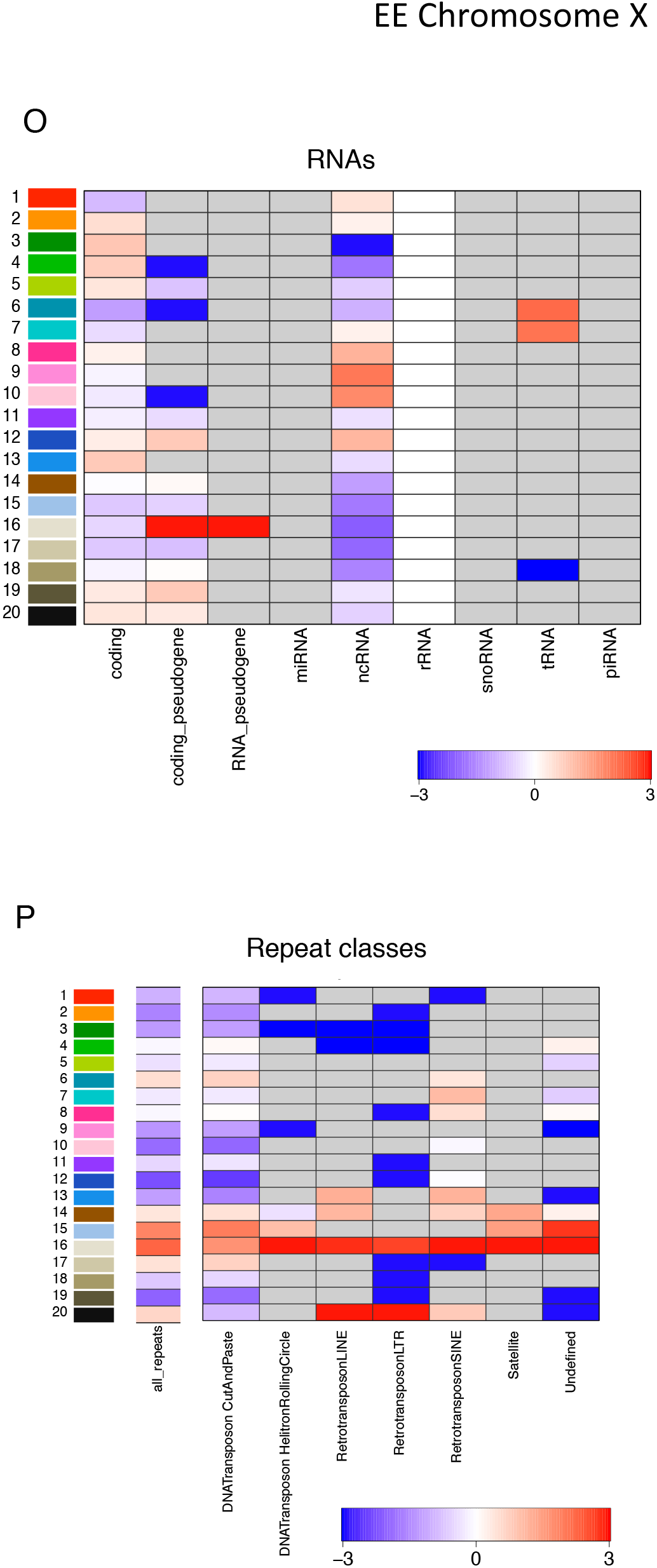

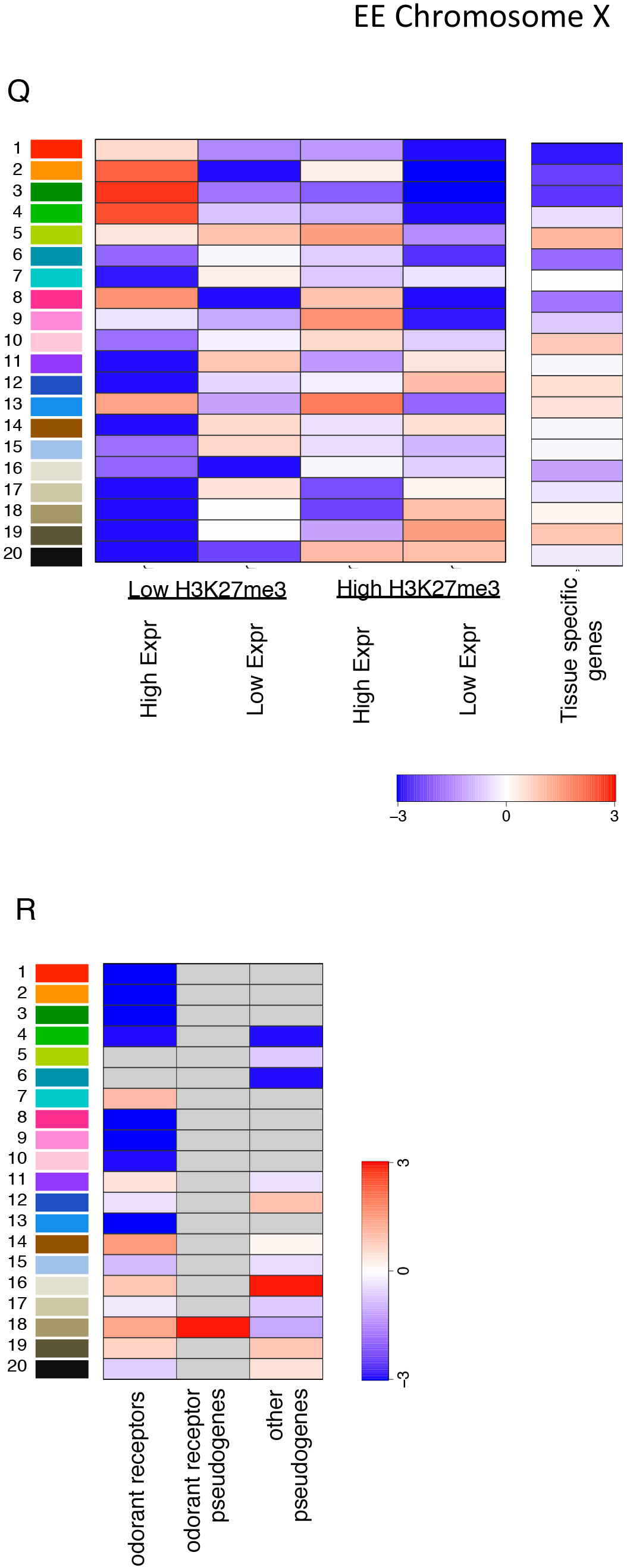

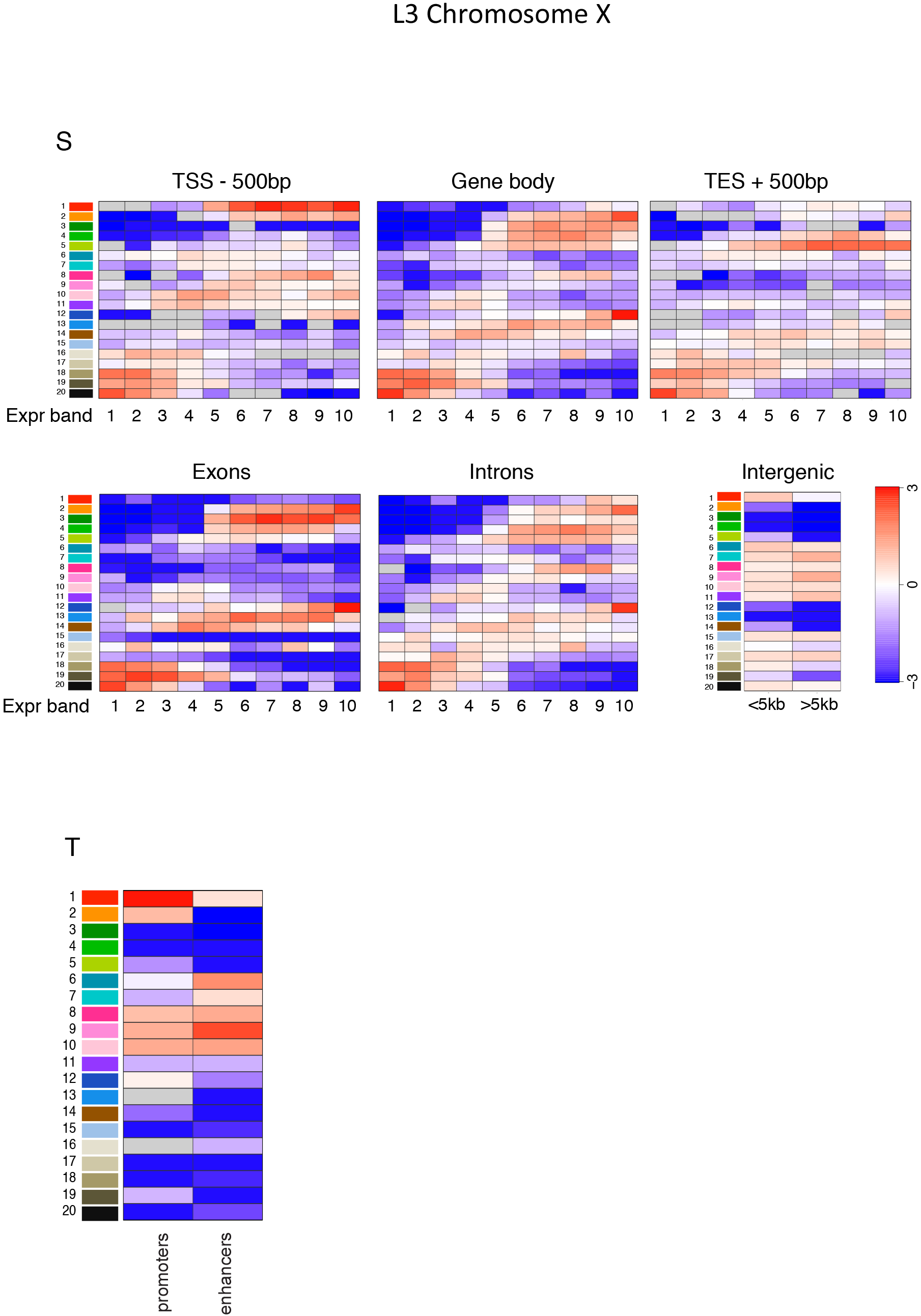

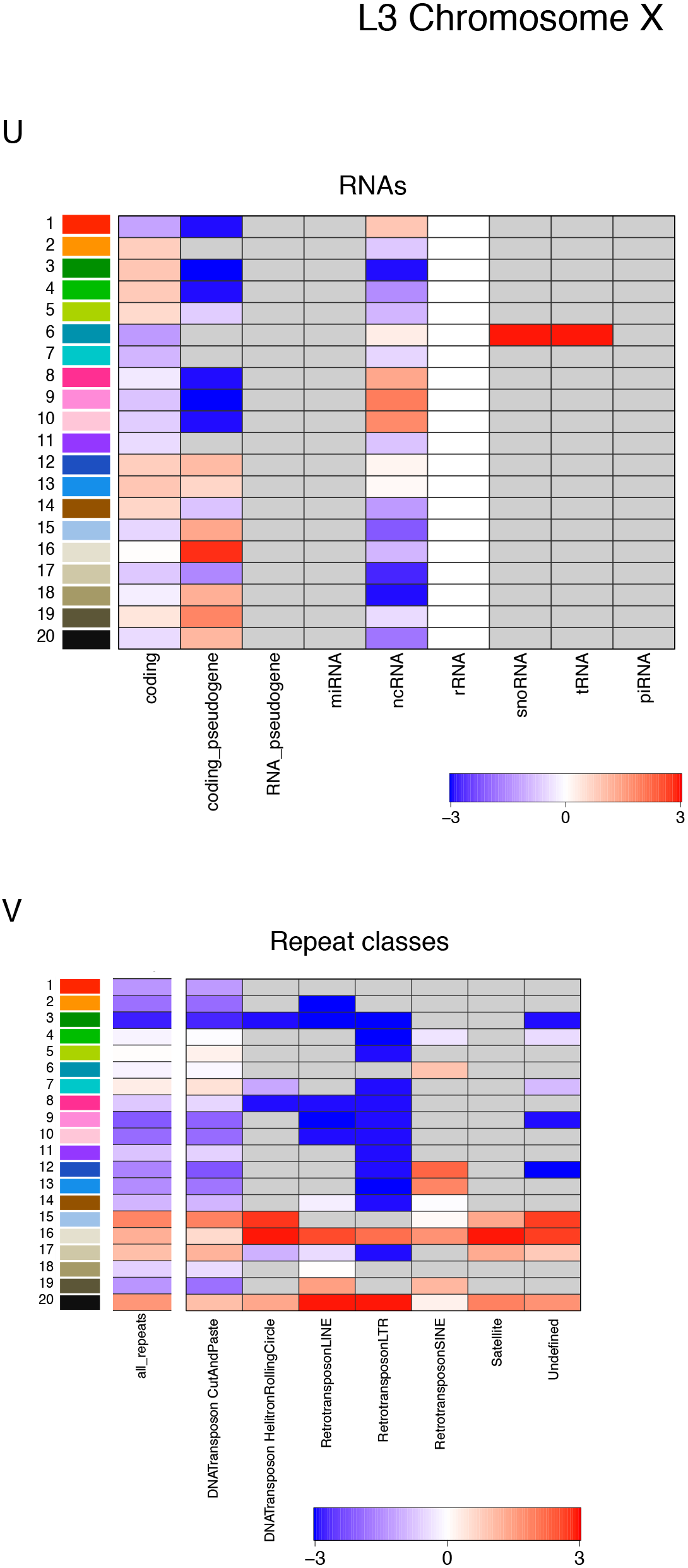

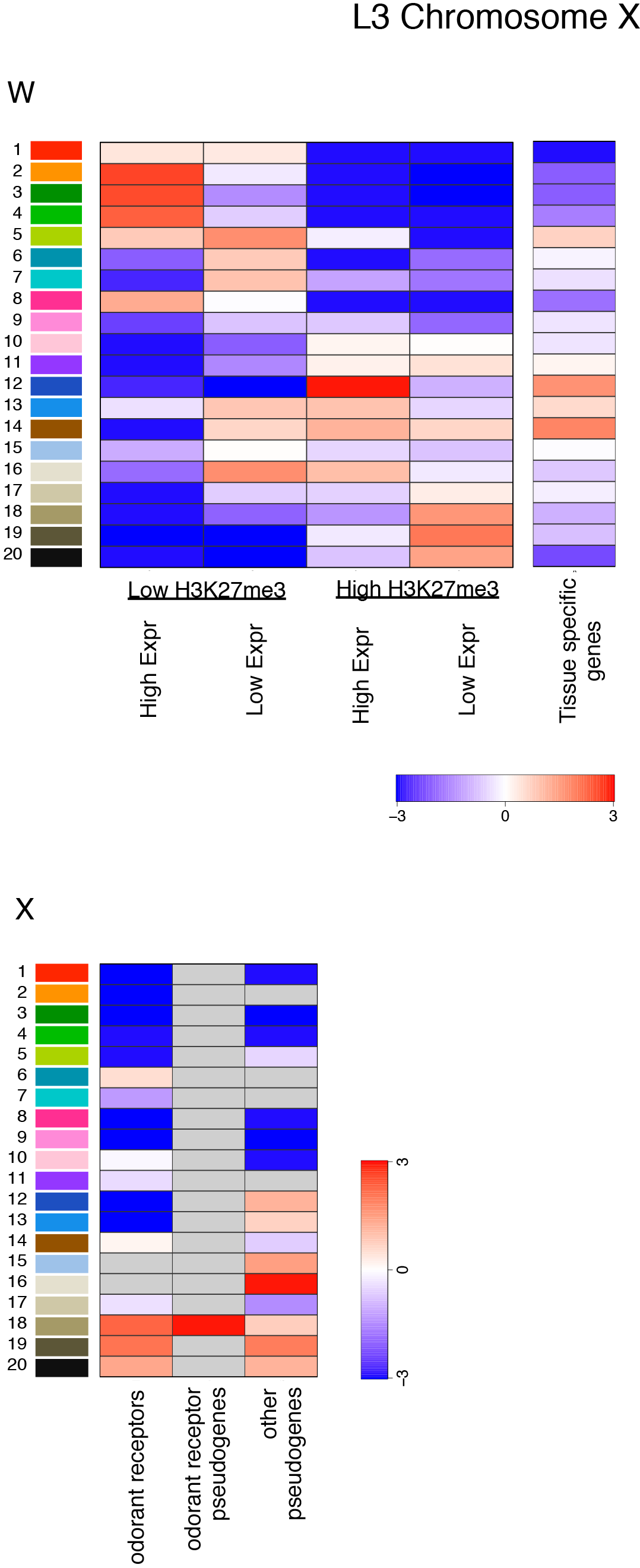

**Figure S5.**
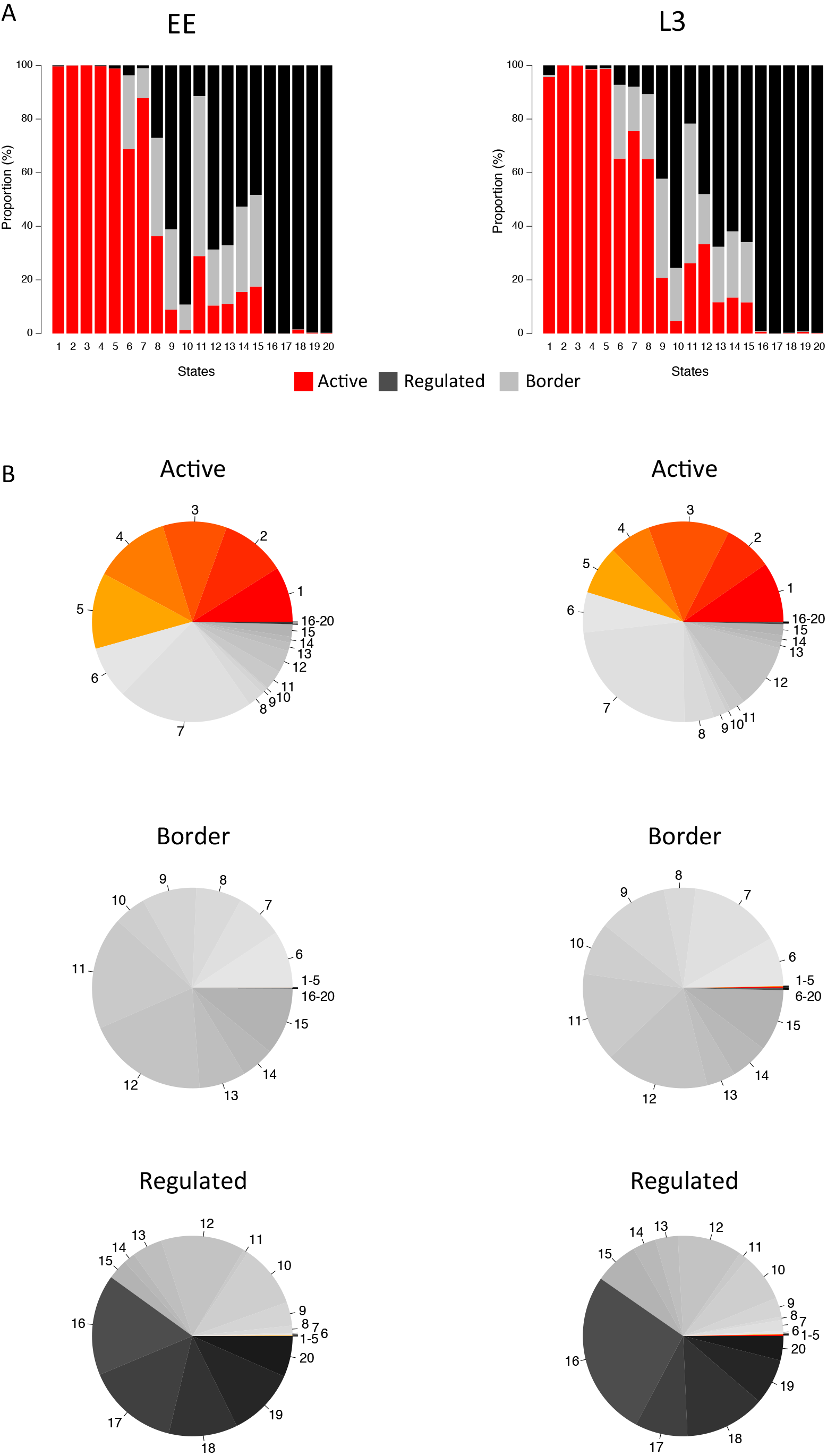
Chromatin state enrichments in different genomic regions and features in L3 autosomes. (A) State enrichments in different genomic features. For each feature, enrichments are shown separately for genes in each decile of expression (expression band 1=lowest). (B) State enrichments in promoters and enhancers. Promoter and enhancer definitions are from (53). (C) State enrichments in coding and different classes of non-coding RNAs. Class definitions are from Wormbase. (D) State enrichments on repeat elements. Repeat charts show state enrichments on all repeat, named classes, and a selection of individual repeats; repeat annotation is from Dfam2.0 (52). (E) State enrichments on groups of genes with low H3K27me3/low gene expression, low H3K27me3/high gene expression, high H3K27me3/ high gene expression, or high H3K27me3/low gene expression, or tissue specific genes. (F) Distribution of states on chromosomes, chromosome regions, odorant receptors, and pseudogenes. Cells are colored grey if there were too few data points for statistical confidence. Scale bars show log2 fold enrichment or underenrichment.

**Figure S6.**
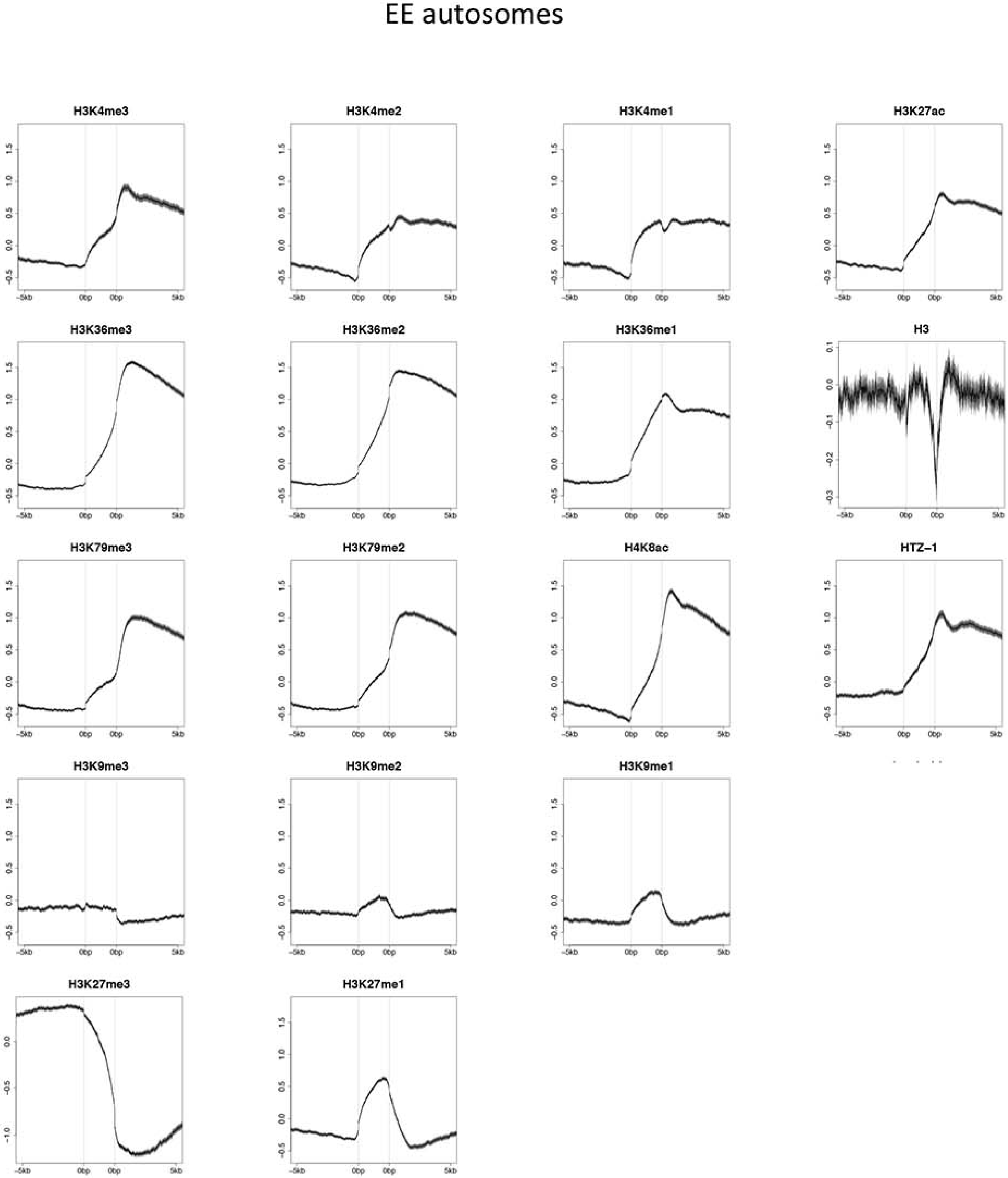
Chromatin state enrichments in different genomic regions and features on EE chromosome X. (A) State enrichments in different genomic features. For each feature, enrichments are shown separately for genes in each decile of expression (expression band 1=lowest). (B) State enrichments in promoters and enhancers. Promoter and enhancer definitions are from (53). (C) State enrichments in coding and different classes of non-coding RNAs. Class definitions are from Wormbase. (D) State enrichments on repeat elements. Repeat charts show state enrichments on all repeats and named classes of repeats. Repeat annotation is from Dfam2.0 (52). (E) State enrichments on groups of genes with low H3K27me3/low gene expression, low H3K27me3/high gene expression, high H3K27me3/ high gene expression, or high H3K27me3/low gene expression, or tissue specific genes. (F) Distribution of states on chromosomes, chromosome regions, odorant receptors, and pseudogenes. Cells are colored grey if there were too few data points for statistical confidence. Scale bars show log2 fold enrichment or underenrichment.

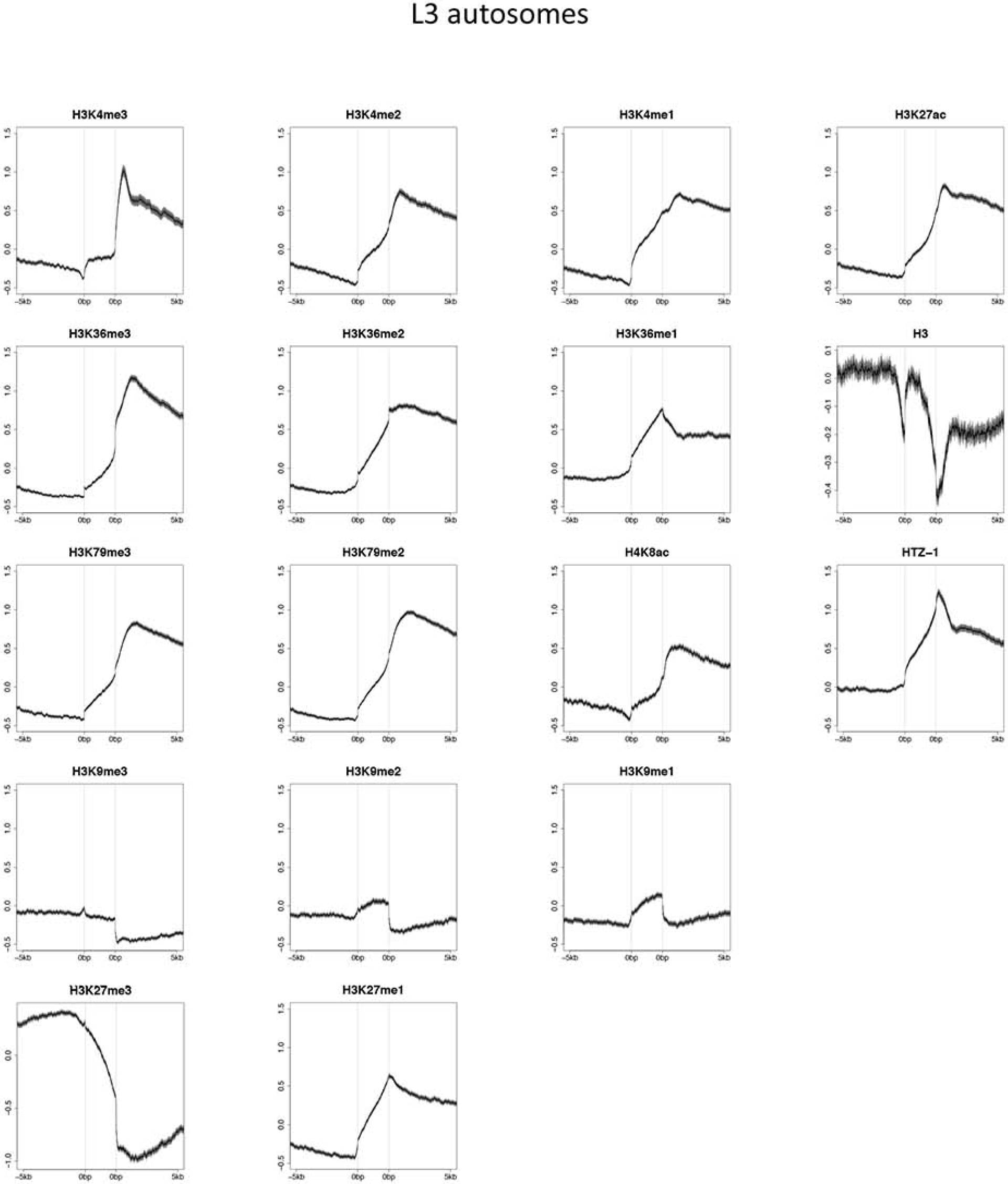

**Figure S7.**
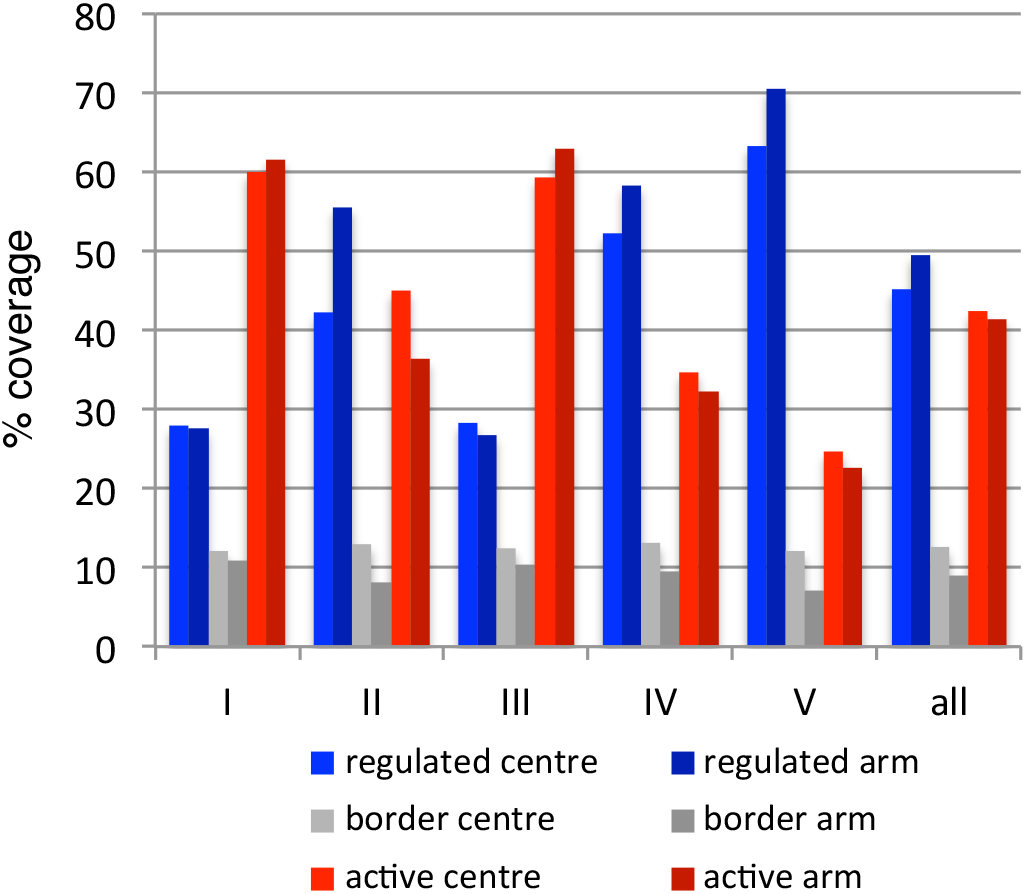
Chromatin state enrichments in different genomic regions and features on L3 chromosome X. (A) State enrichments in different genomic features. For each feature, enrichments are shown separately for genes in each decile of expression (expression band 1=lowest). (B) State enrichments in promoters and enhancers. Promoter and enhancer definitions are from (53). (C) State enrichments in coding and different classes of non-coding RNAs. Class definitions are from Wormbase. (D) State enrichments on repeat elements. Repeat charts show state enrichments on all repeats and named classes of repeats. Repeat annotation is from Dfam2.0 (52). (E) State enrichments on groups of genes with low H3K27me3/low gene expression, low H3K27me3/high gene expression, high H3K27me3/ high gene expression, or high H3K27me3/low gene expression, or tissue specific genes. (F) Distribution of states on chromosomes, chromosome regions, odorant receptors, and pseudogenes. Cells are colored grey if there were too few data points for statistical confidence. Scale bars show log2 fold enrichment or underenrichment.

**Figure S8.**
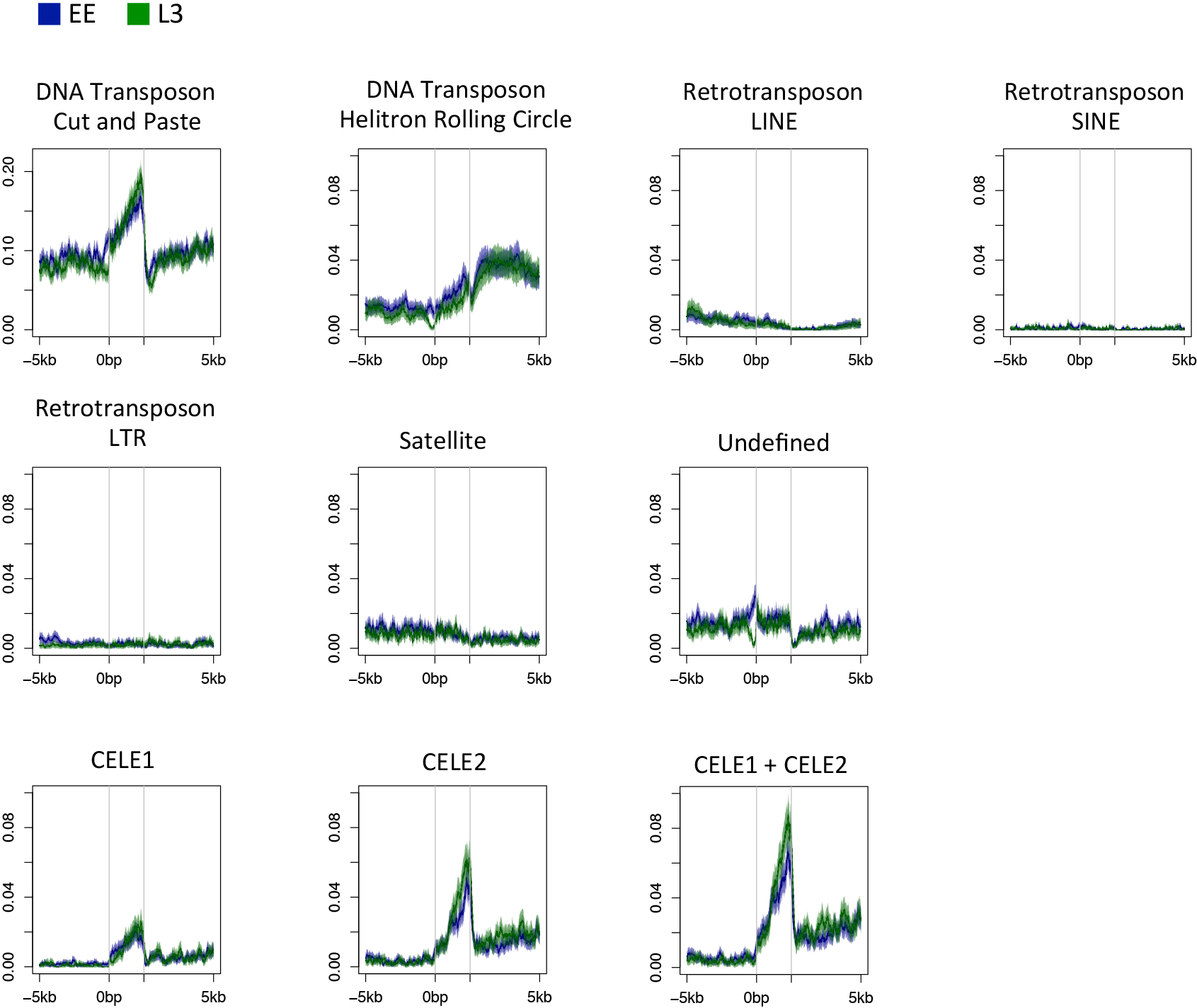
Distribution of chromatin states in EE or L3 active, regulated, and border regions. (A) Proportion of indicated EE or L3 state in active (red), border (grey), and regulated (black) domains. (B) Pie charts showing the distribution of chromatin states in each domain type.

Figure S9. Distribution of histones or histone modifications across EE or L3 active, border, and regulated domains. Plots are centered at borders pseudoscaled at 2.5kb, and show 5 kb into regulated domains (left) and 5kb into active domains (right). Lines show mean signal, darker filled areas show standard error, and lighter filled areas are 95% confidence intervals. Grey vertical lines indicate edges of the border region.

Figure S10. Percent coverage of regulated, border, and active domains in central chromosomal regions and chromosome arms, by chromosome. Within each chromosome, there is a similar proportion of active domains on chromosome arms compared to central regions, and of regulated domains on chromosome arms compared to central regions. However there are between chromosome differences in the overall proportions of active and regulated domains.

Figure S11. Distribution of repeat elements across EE or L3 active, regulated, and border regions. Plots show all repeats, indicated families of repeats, and CELE1 and CELE2 repeats. EE, blue; L3, green. Plots are centered at borders pseudoscaled at 2.5kb, and show 5 kb into regulated domains (left) and 5kb into active domains (right).

Dataset S1. Coordinates of EE and L3 chromatin states.

File of chromosome, start position, end position, and state number. Coordinates are in WS220 and follow BED conventions (start positions are in zero based coordinates and end positions in one based coordinates).

Dataset S2. Coordinates of EE and L3 domains.

Excel file of chromosome, start position, end position of EE and L3 active domains, border regions, and regulated domains (each in separate tab). Additionally, border regions have strand information in column six to indicate if active domain is on the left (-) or on the right (+). Coordinates are in WS220 and follow BED conventions.

**Table S1.** Domain statistics and expected domain sizes and gene numbers

|  | domains (n) | genes (n) | median domain length | expected median domain length | median genes per domain | expected median genes per domain | mean domain length | expected mean domain length | mean genes per domain | expected mean genes per domain |
| --- | --- | --- | --- | --- | --- | --- | --- | --- | --- | --- |
| EE active | 1046 | 6047 | 13054 | 4506 | 4 | 1 | 25318 | 9492 | 5.8 | 2.0 |
| EE border | 2013 | 1914 | 2461 | 600 | 1 | 0 | 5272 | 1865 | 1.0 | 0.4 |
| EE regulated | 1041 | 9424 | 23874 | 13647 | 4 | 2 | 42358 | 21696 | 9.1 | 4.6 |
| L3 active | 1274 | 6198 | 13608 | 4328 | 3 | 1 | 22184 | 8587 | 4.9 | 1.8 |
| L3 border | 2471 | 1996 | 2405 | 500 | 1 | 0 | 4343 | 1794 | 0.8 | 0.4 |
| L3 regulated | 1257 | 9220 | 18670 | 11567 | 3 | 2 | 33619 | 18503 | 7.3 | 3.8 |

**Table S2.** Datasets and Gene expression omnibus accession numbers

| Description | Stage | GEO accession # | Used for chromatin state maps |
| --- | --- | --- | --- |
| H3 | EE | GSE22722 | x |
| H3 | L3 | GSE22734 | x |
| H3K27ac | EE | GSE22748 | x |
| H3K27ac | L3 | GSE25355 | x |
| H3K27me1 | EE | GSE26178 | x |
| H3K27me1 | L3 | GSE26179 | x |
| H3K27me3 | EE | GSE26180 | x |
| H3K27me3 | L3 | GSE22730 | x |
| H3K36me1 | EE | GSE22744 | x |
| H3K36me1 | L3 | GSE26181 | x |
| H3K36me2 | EE | GSE22717 | x |
| H3K36me2 | L3 | GSE22729 | x |
| H3K36me3 | EE | GSE22719 | x |
| H3K36me3 | L3 | GSE22731 | x |
| H3K4me1 | EE | GSE22747 | x |
| H3K4me1 | L3 | GSE25357 | x |
| H3K4me2 | EE | GSE22741 | x |
| H3K4me2 | L3 | GSE22732 | x |
| H3K4me3 | EE | GSE49739 | x |
| H3K4me3 | L3 | GSE28770 | x |
| H3K79me2 | EE | GSE22738 | x |
| H3K79me2 | L3 | GSE26182 | x |
| H3K79me3 | EE | GSE22739 | x |
| H3K79me3 | L3 | GSE26183 | x |
| H3K9me2 | EE | GSE22740 | x |
| H3K9me2 | L3 | GSE22733 | x |
| H3K9me3 | EE | GSE22720 | x |
| H3K9me3 | L3 | GSE22728 | x |
| H4K20me1 | EE | GSE22754 | x |
| H4K20me1 | L3 | GSE25358 | x |
| H4K8ac | EE | GSE26184 | x |
| H4K8ac | L3 | GSE25356 | x |
| HTZ-1 | MxE | GSE50302 | x |
| HTZ-1 | L3 | GSE49717 | x |
| Pol II | EE | GSE25788 |  |
| Pol II | L3 | GSE25792 |  |
| H3K36me3 | EE | GSE38180 |  |
| H3K27me3 | EE | GSE38180 |  |
| MES-4 | EE | GSE38180 |  |
| H3K36me3 | mes-4 RNAi<br>EE | GSE38159 |  |
| H3K27me3 | mes-4 RNAi<br>EE | GSE38159 |  |

